# EZH2 inactivation drives MAPK-dependent vulnerability to MEK inhibition in RAS-mutant CMML

**DOI:** 10.64898/2026.06.02.729524

**Authors:** Eva Gruden, Bianca Perfler, Akshaya Kailasnathan, Panagiota Chaida, Karin Lind, Marie-Christina Mayer, Johannes Foßelteder, Melanie Kienzl, Sayantanee Dutta, Sonja Wurm, Jennifer Neiss, Katarina Vizar Cisarova, Gerald Hoefler, Karl Kashofer, Klaus Geissler, Gerhard Bachmaier, Gudrun Pregartner, Annkristin Heine, Albert Wölfler, Heinz Sill, Andreas Reinisch, Armin Zebisch

## Abstract

Chronic myelomonocytic leukemia (CMML) is a heterogeneous hematologic malignancy with limited therapeutic options. Although *RAS* and *RAS*-modifying mutations (*RAS^mut^*) are common and associated with poor prognosis, targeting *RAS* signaling has shown limited clinical success. Here, we define co-occurrence of *RAS^mut^* and *EZH2* inactivation (*EZH2^inact^*) as a distinct molecular subgroup of CMML characterized by aggressive disease biology. Mechanistically, *RAS^mut^EZH2^inact^* drives selective MAPK/ERK hyperactivation in mature myeloid cells and hematopoietic stem and progenitor compartments. Transcriptomic analysis of a large independent myeloid neoplasm cohort (Beat-AML) further supports this finding, demonstrating selective activation of gene signatures of MAPK/ERK activation in *RAS^mut^EZH2^inact^* cases. Functionally, this signaling activation promotes proliferation and myelomonocytic differentiation in human and murine models. Therapeutically, this MAPK/ERK upregulation confers increased sensitivity to MEK inhibition (MEKi). In a murine *Ras^mut^Ezh2^inact^*-driven CMML model, MEKi suppresses MAPK/ERK activity and reduces leukemic burden by impairing proliferation and myelomonocytic differentiation without inducing cell death. In primary human CMML samples ex vivo, MEKi shows stronger anti-proliferative effects in *RAS^mut^EZH2^inact^* specimens compared with *RAS^mut^* samples. Drug-sensitivity data from the Beat-AML cohort further supported this. Ultimately, these findings were confirmed in a patient-derived xenograft model, where MEKi reduced the leukemic burden by selectively inhibiting the proliferation of transplanted human *RAS^mut^EZH2^inact^* leukemic cells. In summary, our data define *RAS^mut^EZH2^inact^* CMML as a clinically relevant subgroup with selective MAPK/ERK hyperactivation and increased sensitivity to MEKi. They provide a mechanistic explanation for the limited efficacy of MEK inhibitors in unselected CMML and support molecularly guided use of MEK-targeted therapy in this patient population.

## Introduction

Chronic myelomonocytic leukemia (CMML) is an aggressive hematopoietic malignancy with both myelodysplastic (MD) and myeloproliferative (MP) features.^1^ CMML follows an aggressive clinical course with a median overall survival in the higher-risk groups of only about one year.^2, 3^ Allogeneic hematopoietic stem cell transplantation is currently the only curative option.^4^ However, considering that the median age at diagnosis is >70 years, this intensive procedure is not accessible for the majority of CMML patients.^5^ While hypomethylating agents have shown some benefit for MD-CMML, hydroxyurea remains the standard of care for MP-CMML patients.^3, 6^

RAS and RAS-modifying mutations (*RAS^mut^*) are frequently observed in CMML.^7, 8^ They cause a myeloid lineage bias in hematopoietic stem and progenitor cells (HSPC), and promote leukemic proliferation.^9–11^ Indeed, they are mainly associated with MP-CMML.^12–14^ Unlike in many other cancers, *RAS^mut^*in MP-CMML are considered early drivers and dominant clonal events.^14^ Despite their central role in disease biology, *RAS^mut^* are inherently difficult to target therapeutically and have been linked to resistance to targeted therapies in myeloid neoplasms, including the BCL-2 inhibitor venetoclax.^15–17^ Targeting RAS downstream signaling pathways, particularly with MEK inhibitors (MEKi), represents a clinically accessible alternative with a well-established safety profile.^18–20^ However, clinical trials of MEKi in myeloid malignancies, including CMML, have shown limited efficacy, even in cohorts enriched for RAS-mutated disease.^21, 22^

The reason for this limited efficacy remains unclear, but it likely reflects inadequate molecular stratification in the setting of a genetically complex disease.^23–25^ Co-occurring (epi-)genetic events have been shown to influence the oncogenic potential of *RAS^mut^*and the sensitivity to inhibitors targeting *RAS^mut^*-downstream signaling.^26, 27^ We recently demonstrated that *RAS^mut^* often co-occur with the inactivation of the epigenetic modifier EZH2 (*EZH2^inact^*), which leads to transcriptional upregulation of RAS–MAPK/ERK pathway activators and consequent pathway hyperactivation.^28^ In vitro, this genotype results in increased MAPK/ERK signaling and confers preferential sensitivity to pharmacological MEKi, suggesting a context-dependent therapeutic vulnerability. However, these findings were derived from in vitro studies and lacked in vivo and translational validation, limiting direct clinical applicability.

Here, we comprehensively characterize *RAS^mut^EZH2^inact^*CMML in a translational setting, including transgenic mice, primary patient samples, and a patient-derived xenograft model. We demonstrate that this genotype drives selective amplification of RAS signaling via the MAPK/ERK pathway, resulting in pronounced sensitivity to pharmacologic MEKi. These findings provide a mechanistic explanation for the limited efficacy of MEK inhibitors in unselected CMML populations and suggest that effective clinical application will require molecularly guided patient selection that accounts for co-occurring genetic and epigenetic alterations, such as RAS mutations and EZH2 inactivation.

## Materials and Methods

### Mice and patient-derived xenografts

Transgenic mice were acquired from The Jackson Laboratory (www.jax.org/) on a C57BL/6 background. Mice carrying *Nras^G12D^* (Nras C57BL/6-Nrastm1Tyj/J), *Ezh2* deletion (C57BL/6-Ezh2tm2Sho/J), and the Mx1-Cre promoter (C57BL/6-Tg(Mx1-cre)1Cgn/J) were intercrossed to generate *Mx1-Cre*-*Nras^G12D/Wt^* (hereafter referred as *Nras^G12D^), Mx1-Cre*-*Nras^G12D/Wt^Ezh2^−/−^*(*Nras^G12D^Ezh2^−/−^), Mx1-Cre*-*Nras^G12D/Wt^Ezh2^+/−^* (*Nras^G12D^Ezh2^+/−^)* and *Nras/Ezh2* wildtype mice (*Wt).* Recombination was induced by polyinosinic-polycytidylic acid (pIpC) injection at 5-6 weeks of age. For xenotransplantation studies, twelve– to thirteen-week-old female NSG mice (Charles River Laboratories, Wilmington, MA) were used. Approximately 2×10^6^ cells from patient LB-MUG-294 were transplanted intrafemorally into sub-lethally irradiated (1 Gy) recipients. Engraftment was verified 8 weeks after transplantation. All mice were maintained under specific pathogen-free conditions. Experiments were approved by the Austrian Federal Ministry of Science and Research (BMBWF-66.010/0190-V/3b/2018). Treatment schedules and detailed experimental procedures are provided in the Supplemental Materials.

### Primary human samples

Human bone marrow (BM) and peripheral blood samples were collected at Medical University of Graz (provided by the Biobank of Medical University of Graz, Cohort 6002_14, Myeloid Neoplasias Collection) as described previously.^29–32^ Sample collection and use were approved by the local institutional review board (approval numbers 30-464ex17/18 and 1212/2024) and performed in accordance with the Declaration of Helsinki. Additional information about biobanking, clinical and molecular patient characteristics, and detailed information about ex vivo experiments are provided in Supplemental Table 1.

### Whole transcriptome RNA sequencing (RNA-Seq)

RNA-seq from transgenic *Nras^G12D^Ezh2^−/−^* mice after Mirdametinib treatment was conducted as a service by Azenta Life Sciences (Griesheim, Germany), and raw reads in FASTQ format were provided. Data analysis and gene set enrichment analysis were performed as described previously.^33^ All raw data are publicly available via the NCBI Gene Expression Omnibus (accession number will be provided upon publication).

### Database retrieval and statistical analysis

Overall and AML-free survival for CMML patients were obtained from the MDS-IWG_IPSS-M cohort^34^ via cBioPortal (https://www.cbioportal.org/)^35, 36^ on 01-AUG-2025. Ex vivo drug response data and the corresponding *RAS^mut^*and *EZH2^inact^* status for AML were obtained from the Beat-AML cohort (downloaded 24-APR-2025 and extracted from Supplemental Table S10).^37^ Canonical pathway gene sets (BIOCARTA_ERK_PATHWAY, BIOCARTA_AKT_PATHWAY and BIOCARTA_p38/MAPK_PATHWAY) were downloaded from https://www.gsea-msigdb.org/gsea/index.jsp.^38, 39^ Additional details about database analyses are provided in the Supplemental Materials.

Statistical analyses were performed using GraphPad Prism 8.0.2 or 10.0.3 (GraphPad Software, Boston, MA) as well as R version 4.5.0. Experimental group comparisons were performed from at least three independent repetitions and analyzed by *t*-tests, Wilcoxon matched-pairs signed-rank test or ANOVA, as depicted in the respective figure legends. In human patients, dichotomous variables were compared by Fisher’s exact and continuous variables by Mann Whitney U test. We performed Kaplan-Meier survival analyses and compared groups using the log-rank test. In all cases, the results are considered exploratory and a p value < 0.05 was considered statistically significant.

## Results

### *RAS^mut^EZH2^inact^* co-occurrence is associated with MP-CMML and an aggressive clinical course

To assess the clinical impact of *RAS^mut^EZH2^inact^*co-occurrence, we analyzed a publicly available dataset of 399 CMML patients^34^ and grouped patients into *RAS^wt^*, *RAS^mut^* and *RAS^mut^EZH2^inact^*. Due to the absence of available transcriptomic data for this cohort, *EZH2^inact^* was defined as either hetero– or homozygous mutation of *EZH2* and/or del(7q)/-7, based on the localization of EZH2 on the long arm of chromosome 7. This selection was based on data showing that both biallelic (i.e. by homozygous mutations) and monoallelic (i.e. by heterozygous mutations and del[7q] or –7) alterations in *EZH2* are associated with adverse clinical outcomes, decreased EZH2 expression, and activation of EZH2 downstream targets.^40–42^ Patients harboring *RAS^mut^* and *EZH2^inact^* showed a significantly shorter overall (OS, Figure 1A) and AML-free survival (Figure 1B) compared to both *RAS^wt^* and *RAS^mut^*groups.

**Figure 1:**
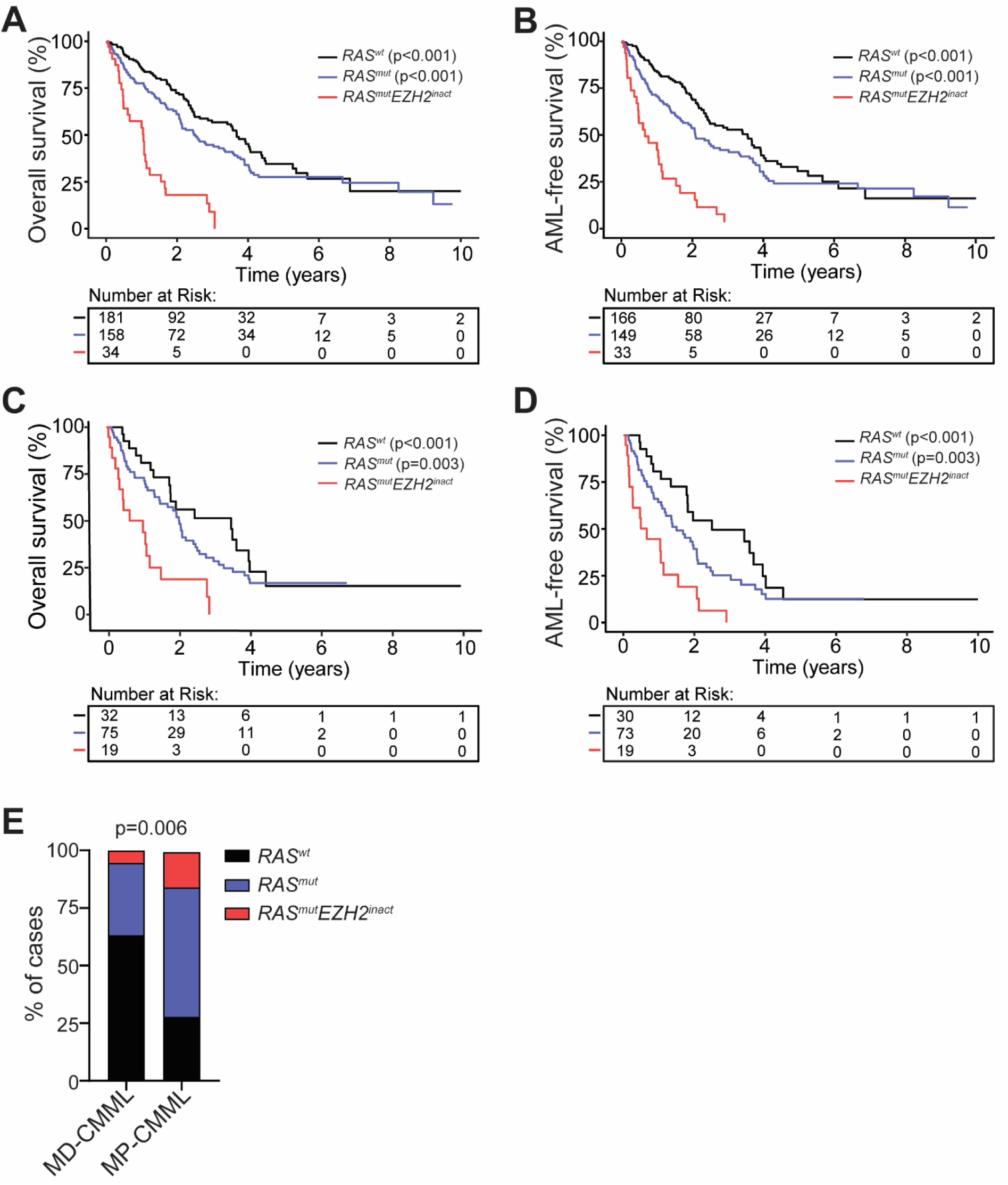
*RAS^mut^EZH2^inact^* co-occurrence is associated with MP-CMML and an aggressive clinical course. (A-B) Overall survival (OS, A) and AML-free survival (B) of CMML patients from the IPSS-M cohort.^34^ Patients were grouped into *RAS^wt^*, *RAS^mut^* or *RAS^mut^EZH2^inact^*. *RAS^mut^EZH2^inact^* co-occurrence was associated with a shortened OS (median 13 vs. 30 or 44 months) and AML-free survival (median 8 vs. 25 or 43 months) compared to *RAS^mut^* or *RAS^wt^* groups. (C-D) OS (C) and AML-free survival (D) in MP-CMML patients, showing inferior outcomes of *RAS^mut^EZH2^inact^*patients compared to *RAS^mut^* or *RAS^wt^* cases in this subgroup as well (median OS 12 vs 25 or 42 months; median AML-free survival 8 vs 18 or 30 months). Kaplan-Meier survival curves were compared using the log-rank test, followed by pairwise comparisons with Holm correction for multiple testing, only comparisons versus *RAS^mut^EZH2^inact^* are shown. (E) Distribution of *RAS^wt^*, *RAS^mut^* and *RAS^mut^EZH2^inac^*^t^ across MD-CMML and MP-CMML. *RAS^mut^EZH2^inact^*were enriched in MP-CMML compared to MD-CMML (15% vs 5%, p=0.006 by Fisheŕs exact test.

As RAS mutations are associated with the more aggressive MP-CMML subtype,^12–14^ we next limited our analyses to cases with MP-CMML. Again, *RAS^mut^EZH2^inact^* patients had shorter OS (Figure 1C) and AML-free survival (Figure 1D) compared to both *RAS^wt^* and *RAS^mut^* groups. Of note, similar to *RAS^mut^* patients,^12–14^ the frequency of *RAS^mut^EZH2^inact^*was increased in MP-CMML as well (Figure 1E).

### *Nras^G12D^Ezh2^−/−^* mice develop an aggressive MP-CMML-like disease with selective activation of the RAS-MAPK/ERK pathway

To further investigate the pathological features and underlying mechanisms associated with the co-occurrence of RAS mutations and EZH2 inactivation in vivo, we employed a mouse model of chronic myeloproliferation that resembles MP-CMML, in which disease development is driven by the *Nras^G12D^Ezh2^−/−^*genotype.^43^ In this model both heterozygous oncogenic Ras mutation (*Nras^G12D+/−^*) and deletion of Ezh2 (Ezh2^−/−^) are driven by an Mx1-Cre promoter (Figure 2A). *Nras^G12D^Ezh2^−/−^*mice developed a highly aggressive disease phenotype characterized by shorter survival (Figure 2B) compared to all other genotypes, including mice carrying *Nras^G12D^*only (median survival 115 vs 366 days). They also demonstrated splenomegaly (Figure 2C), as well as increased white blood cell (WBC) and platelet counts (Figure 2D). Flow cytometric analysis revealed a marked expansion of the myelomonocytic cell population (CD11b^+^Gr1^+^ cells) in PB (Figure 2E), spleen (Figure 2F) and BM (Figure 2G) of *Nras^G12D^Ezh2^−/−^*mice.

**Figure 2:**
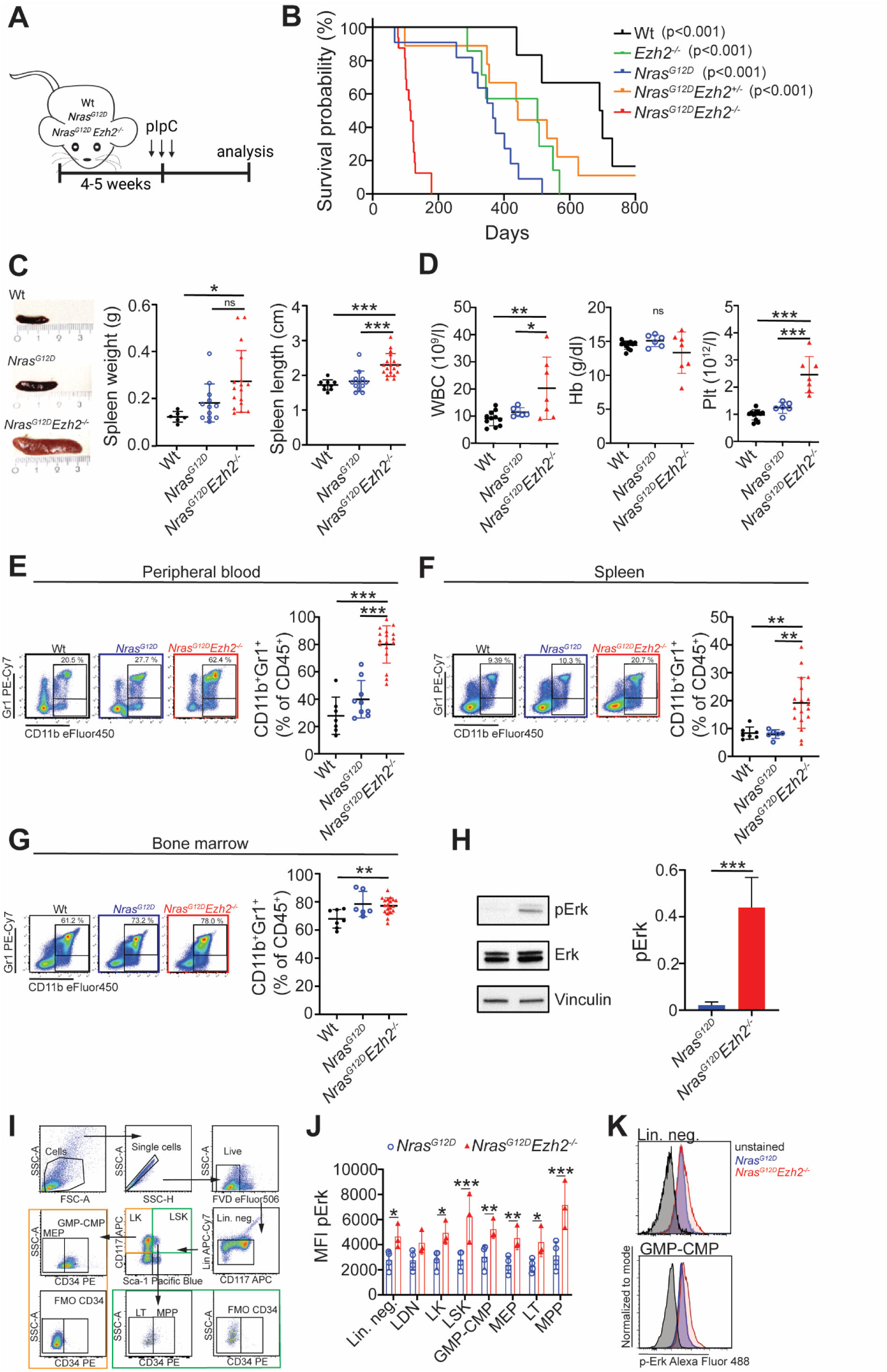
*Nras^G12D^Ezh2^−/−^* mice develop an aggressive CMML-like disease with activation of the RAS-MAPK/ERK pathway. (A) Schematic representation of the experimental design. (B) Overall survival of mice with indicated genotypes (n=6-16). Kaplan-Meier survival curves were compared using the log-rank test, followed by pairwise comparisons mice with Holm-Sidak correction for multiple testing; only comparisons versus *Nras^G12D^Ezh2^−/−^* mice are shown. (C) Representative images and quantification of spleen weight and length of Wt, *Nras^G12D^* and *Nras^G12D^Ezh2^−/−^*mice. (D) Peripheral blood counts, including white blood cells (WBC), hemoglobin (Hb) and platelets (Plt), measured 15 weeks post-pIpC or when moribund. (E-G) Representative pseudocolor plots and quantification of myelomonocytic (CD11b^+^Gr1^+^) cells in peripheral blood (E), spleen (F), and BM (G) of mice 6 months post-pIpC or when moribund. Data were analyzed by one-way ANOVA (n=6-16) using Tukey’s correction for pairwise comparisons. (H) Representative Immunoblot and quantification of pErk levels in total BM of *Nras^G12D^* and *Nras^G12D^Ezh2^−/−^*mice. Comparisons were performed from three independent experiments using an unpaired *t* test. (I-K) Phospho-flow analysis of lineage depleted BM. (I) Representative gating strategy for HSPC compartments evaluated for pErk. (J) Mean fluorescence intensity (MFI) of pErk across progenitor compartments. (K) Representative histogram overlays of pErk staining in *Nras^G12D^* and *Nras^G12D^Ezh2^−/−^* total lineage negative cells (top) or in the GMP-CMP cell compartment (bottom). n=3-4; analyzed with multiple *t* tests. Wt, wild-type; pIpC, polyinosinic:polycytidylic acid; FVD, fixable viability dye; Lin.neg., lineage negative cells; LDN, (Lin^-^, Sca-1^-^, CD117^-^ cells); LK, (Lin^-^, Sca-1^-^, CD117^+^ cells); LSK, (Lin^-^, Sca-1^+^, CD117^+^ cells); GMP-CMP, Granulocyte Macrophage and Common Myeloid progenitors (Lin^-^ Sca-1^-^ c-kit^+^ CD34^+^); MEP, Megakaryocyte Erythroid Progenitors (Lin^-^ Sca1^-^ c-kit^+^ CD34^-^); LT, long term hematopoietic stem cells (Lin^-^ Sca-1^+^ c-kit^+^ CD34^-^), MPP, multipotent progenitor (Lin^-^ Sca-1^+^ c-kit^+^ CD34^+^); ns, not significant. Data are shown as mean ± SD; *p < .05, **p < .01, ***p < .001.

As we have previously shown a hyperactivation of the RAS-MAPK/ERK pathway in *RAS^mut^* myeloid cell lines with additional EZH2 knockdown or pharmacologic inhibition of EZH2,^28^ we next assessed pathway activation in vivo. Immunoblot analysis of total BM cells demonstrated significantly increased Erk phosphorylation (pErk) in *Nras^G12D^Ezh2^−/−^*mice compared to *Nras^G12D^* animals (Figure 2H). Phospho-flow analysis of lineage-depleted BM cells further confirmed these results and demonstrated increased pERK levels in total lineage-negative cells and in the majority of HSPC compartments such as LK, LSK, GMP-CMP, MEP, LT, and MPP (Figure 2I-K). Importantly, amplification of RAS-downstream signaling in the *Nras^G12D^Ezh2^−/−^* genotype was selectively restricted to the MAPK/ERK pathway, whereas other RAS-downstream cascades, including p38/MAPK and PI3K/AKT, remained unaffected (Supplemental Fig. 1). Of note, complete Ezh2 deletion was required to induce both the aggressive disease phenotype and MAPK/ERK hyperactivation, since *Nras^G12D^Ezh2^+/−^* mice displayed a milder disease course (Supplemental Figure 2A-B), only slightly decreased Ezh2 expression, and absence of MAPK hyperactivation (Supplemental Figure 2C). This is in contrast to human disease, where both monoallelic and biallelic EZH2 inactivation was associated with adverse clinical outcomes, decreased EZH2 expression, and activation of EZH2 downstream targets.^40–42^ Therefore, subsequent experiments were focused to *Nras^G12D^Ezh2^−/−^*mice only.

### MEK inhibition suppresses RAS-MAPK/ERK signaling and mediates a pronounced anti-leukemic effect in *Nras^G12D^Ezh2^−/−^* mice

Given the observed MAPK/ERK hyperactivation in *Nras^G12D^Ezh2^−/−^* mice, we hypothesized that this genotype would exhibit increased therapeutic sensitivity to MEKi. Therefore, we treated these mice with the U.S. Food and Drug Administration (FDA)– and European Medicines Agency (EMA)-approved MEK inhibitor Mirdametinib (1.5 mg/kg, p.o.)^44^ or vehicle for 10 weeks (Figure 3A). MEKi significantly reduced pErk levels in HSPC compartments in the BM (Figure 3B). This was associated with a marked anti-leukemic effect, as evidenced by significantly prolonged survival (Figure 3C), reduced splenomegaly (Figure 3D-F), and decreased myelomonocytic infiltration of the spleen (Figure 3G). PB analysis was performed every second week during the treatment period and revealed a progressive increase in WBC counts in vehicle-treated mice, whereas WBC levels remained stable in Mirdametinib-treated animals (Figure 3H). In contrast, hemoglobin and platelet counts were not significantly altered in either group over the treatment period (Figure 3I-J). Importantly, differential blood counts revealed that Mirdametinib treatment reversed the myelomonocytic bias observed in vehicle-treated mice (Figure 3K). This was further confirmed via flow cytometry, showing a significant reduction of the CD11b^+^ Gr1^+^ population in the Mirdametinib-treated mice (Figure 3L-M).

**Figure 3:**
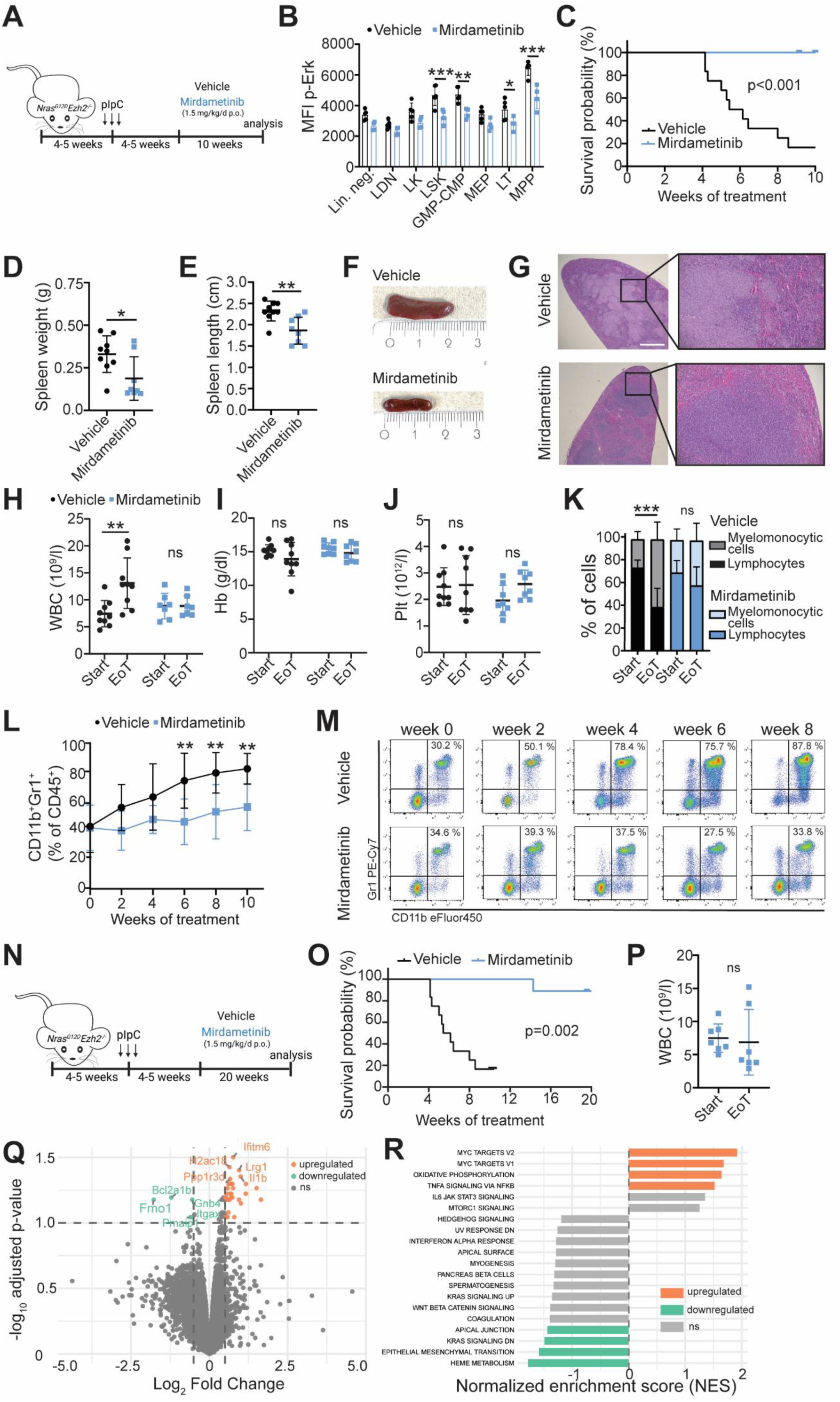
MEK inhibition with Mirdametinib suppresses RAS-MAPK/ERK signaling and mediates a pronounced anti-leukemic effect in *Nras^G12D^Ezh2^−/−^* mice. (A) Schematic overview of the short-term Mirdametinib MEK inhibitor (MEKi) treatment design. Mice were sacrificed after 10 weeks of treatment or when moribund. (B) Phospho-flow analysis of lineage depleted BM (gating strategy in Fig. 2I) from mice treated either with vehicle or Mirdametinib for 10 days. n=4-5; analyzed with multiple *t* tests. (C) Survival of mice after the start of vehicle or Mirdametinib treatment. Kaplan-Meier analysis was performed groups were compared using the log-rank test (n=11-12). (D-F) Spleen weight (D), spleen length (E) and representative spleen images (F) at the experimental end point. n=8-9; analyzed with unpaired *t* test. (G) Representative hematoxylin & eosin staining of spleen sections from vehicle (top) or Mirdametinib-treated (bottom) animals. The scale bar represents 100 μm. (H-J) Peripheral blood counts, including white blood cells (WBC, H), hemoglobin (Hb, I) and platelets (Plt, J) comparing start and end of treatment. (K) Myelomonocytic bias, assessed by differential WBC counts (relative proportions of lymphocytes and myelomonocytic cells), at start and end of treatment. n=7-9; analyzed by two-way ANOVA followed by Sidak’s multiple comparisons test. (L-M) Quantification (L) and representative pseudocolor plots (M) of CD11b^+^Gr1^+^ myelomonocytic cells in peripheral blood during treatment n=8-9; analyzed with two-way ANOVA, using Sidak’s correction for multiple comparisons comparing vehicle to Trametinib treatment. (N) Schematic overview of the long-term Mirdametinib MEKi treatment design. (O) Survival of mice in the long-term cohort after the start of vehicle or Mirdametinib treatment. Kaplan-Meier analysis was performed and groups were compared using the log-rank test (n=9-12). (P) Peripheral blood WBC counts of mice treated long-term with Mirdametinib comparing start and end of treatment. n=7; analyzed by paired *t* test. (Q-R) Whole transcriptome analysis of total BM cells of mice treated with vehicle or Mirdametinib for 1 week. (Q) Volcano plot of differentially expressed genes (FDR<0.05). (R) Gene Set Enrichment analysis (GSEA) comparing Mirdametinib (n=4) and vehicle-treated (n=3) animals. pIpC, polyinosinic:polycytidylic acid; Lin.neg., lineage negative cells; LDN, (Lin^-^, Sca-1^-^, CD117^-^cells); LK, (Lin^-^, Sca-1^-^, CD117^+^ cells); LSK, (Lin^-^, Sca-1^+^, CD117^+^ cells); GMP-CMP, Granulocyte Macrophage and Common Myeloid progenitors (Lin^-^ Sca-1^-^ c-kit^+^ CD34^+^); MEP, Megakaryocyte Erythroid Progenitors (Lin^-^ Sca1^-^ c-kit^+^ CD34^-^); LT, long term hematopoietic stem cells (Lin^-^ Sca-1^+^ c-kit^+^ CD34^-^), MPP, multipotent progenitor (Lin^-^ Sca-1^+^ c-kit^+^ CD34^+^); EoT, end of treatment; MMC, myelomonocytic cells; ns, not significant. Data are shown as mean ± SD; *p < .05, **p < .01, ***p < .001.

Long-term treatment of more than 5 months with Mirdametinib resulted in sustained disease control and significantly prolonged survival, without inducing cytopenias or other toxicities (Figure 3N–P). To further explore the underlying mechanism, we performed bulk RNASeq analysis of total BM cells derived from *Nras^G12D^Ezh2^−/−^* mice treated with Mirdametinib or vehicle. These analyses revealed a suppression of oncogenic signaling and mesenchymal programs as well as a compensatory increase in metabolic and stress responses (Figure 3Q-R).

To further validate these findings, we treated *Nras^G12D^Ezh2^−/−^* mice with Trametinib, another FDA– and EMA-approved MEK inhibitor (Figure 4A). Compared to Mirdametinib, Trametinib exhibits higher cellular potency.^45^ Indeed, Trametinib treatment reproduced the anti-leukemic effects at approximately threefold lower doses compared to Mirdametinib and resulted in significantly prolonged survival (Figure 4B) and reduced splenomegaly (Figure 4C-E). Moreover, it reduced the myelomonocytic bias (Figure 4F), improved hematologic parameters (Figure 4G-I), and decreased the percentage of CD11b^+^Gr1^+^ cells in PB (Figure 4J).

**Figure 4:**
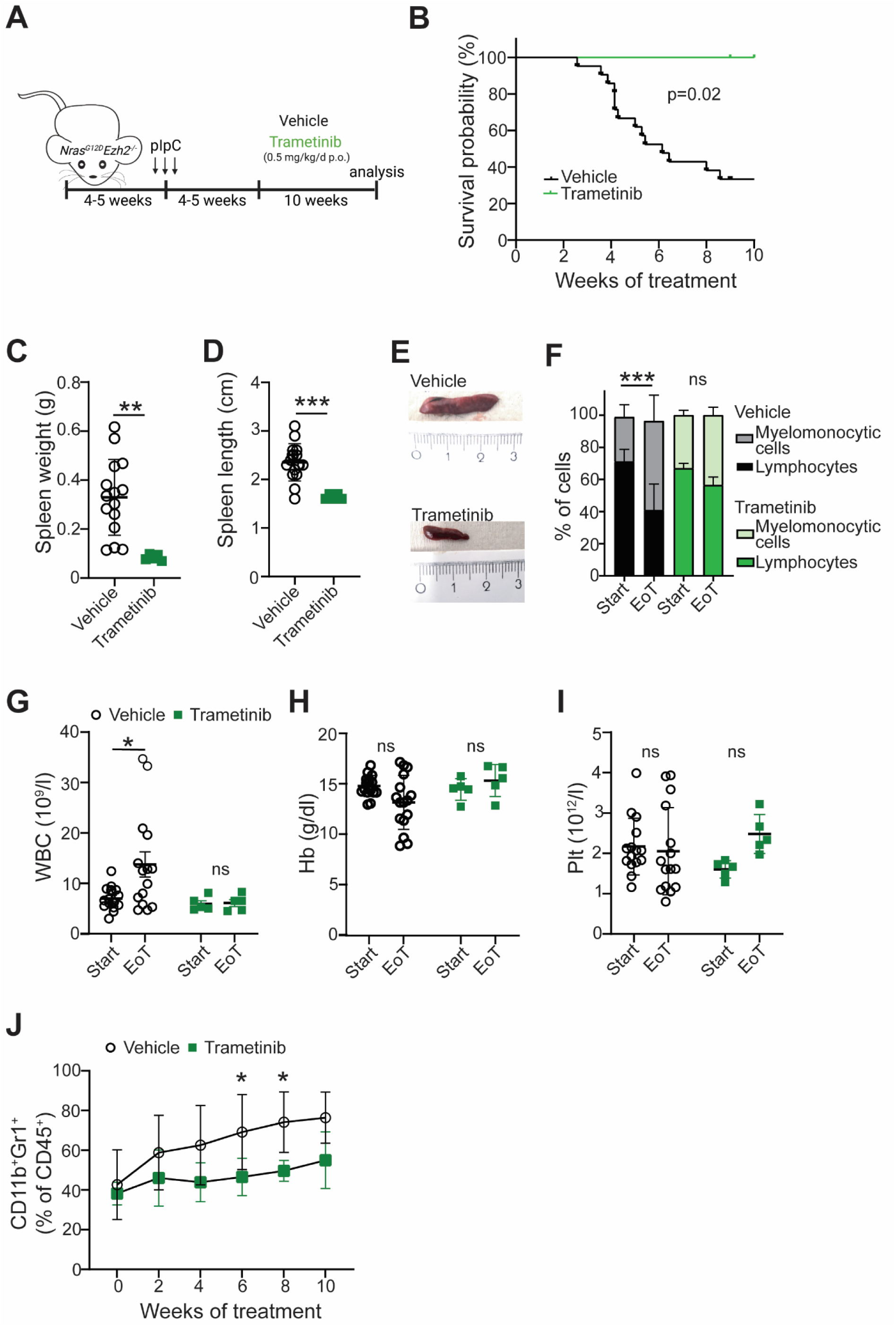
MEK inhibition with Trametinib induces a pronounced anti-leukemic effect in *Nras^G12D^Ezh2^−/−^* mice. (A) Schematic overview of the Trametinib MEK inhibitor (MEKi) treatment design. Mice were sacrificed after 10 weeks of treatment or moribund. (B) Survival of mice after the start of vehicle or Trametinib treatment. Kaplan-Meier analysis was performed and groups were compared using the log-rank test (n=5-21). (C-E) Spleen weight (C), length (D), and representative spleen images (E) of vehicle– and Trametinib-treated mice at the experimental end point. n=5-15; analyzed with unpaired *t* test. (F) Myelomonocytic bias, assessed by differential WBC counts (relative proportions of lymphocytes and myelomonocytic cells), at start and end of treatment. n=5-15; analyzed by two-way ANOVA using Sidak’s correction for multiple comparisons. (G-I) Peripheral blood counts of white blood cells (WBC, G), platelets (Plt, H) and hemoglobin (Hb, I), comparing start and end of treatment. n=5-18; analyzed by two-way ANOVA using Sidak’s correction for multiple comparisons. (J) Quantification of CD11b^+^Gr1^+^ myelomonocytic cells in peripheral blood during treatment n=5-15; analyzed with two-way ANOVA, using Sidak’s correction for multiple comparisons, comparing vehicle to Trametinib treatment. pIpC, polyinosinic:polycytidylic acid; EoT, end of treatment; MMC, myelomonocytic cells; ns, not significant. Data are shown as mean ± SD; *p < .05, **p < .01, ***p < .001.

### MEKi suppresses *Nras^G12D^Ezh2^−/−^*-driven myelomonocytic lineage commitment and proliferation without inducing cell death

Since MEK inhibition reduced the myelomonocytic bias of *Nras^G12D^Ezh2^−/−^* in vivo, we assessed whether MEKi affects proliferation or cell survival. Ex vivo EdU-based cell-cycle assays (Figure 5A) demonstrated that both MEK inhibitors dose-dependently and significantly reduced the percentage of cells in S-phase in *Nras^G12D^Ezh2^−/−^* CD11b^+^ BM cells (Figure 5B-C). In contrast, Annexin V/7-AAD staining revealed no significant induction of cell death at the tested concentrations (Figure 5D-E). In addition to its effects on proliferation, we assessed the impact of MEKi on myelomonocytic lineage commitment. Treatment of lineage-depleted *Nras^G12D^Ezh2^−/−^* BM cells with Trametinib resulted in a reduced expansion of CD11b^+^Gr1^+^ cells over time under stimulation with low-dose granulocyte-macrophage colony-stimulating factor (GM-CSF) (Figure 5F-H).

**Figure 5:**
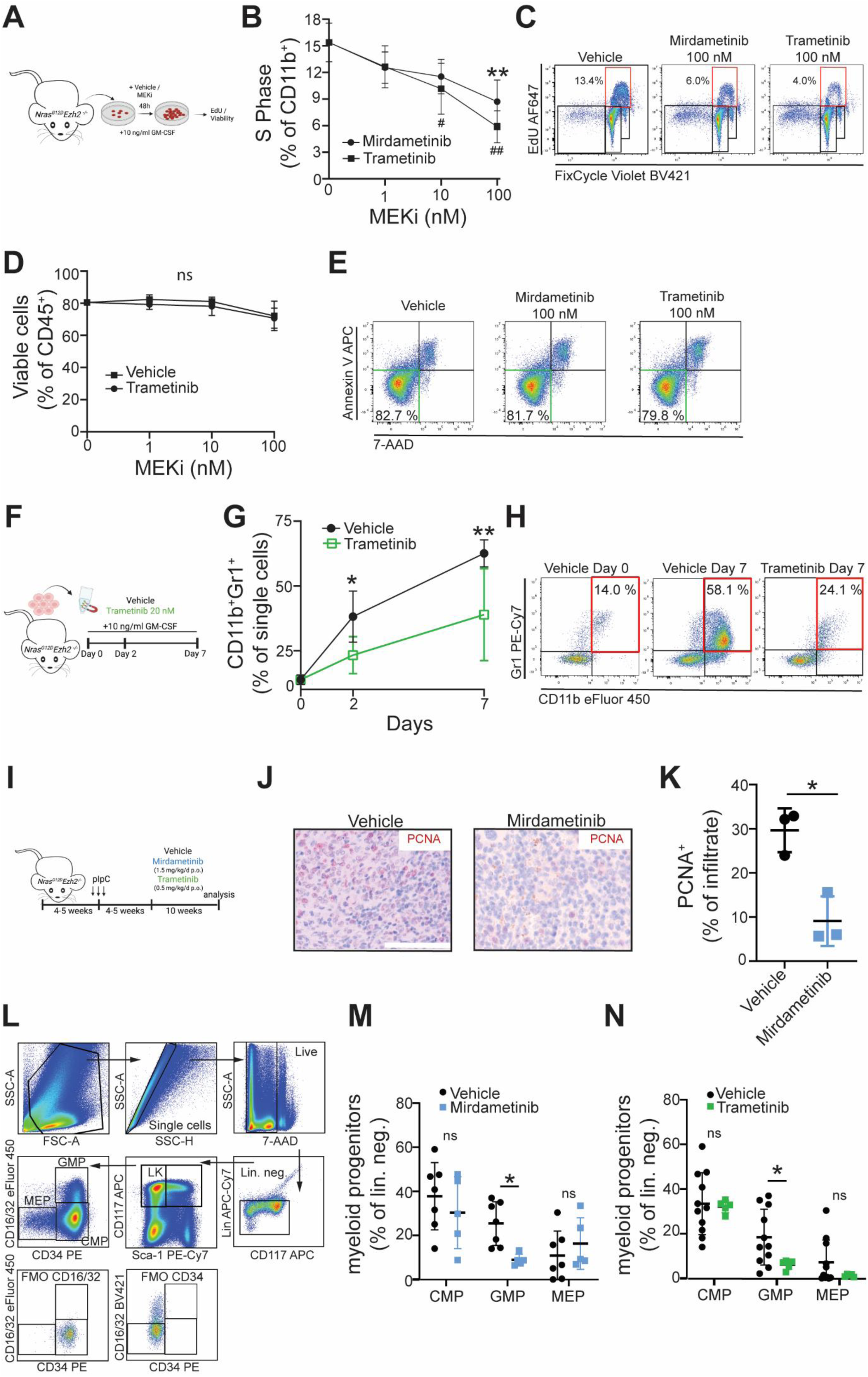
MEK inhibition suppresses *Nras^G12D^Ezh2^−/−^*-driven myelomonocytic lineage commitment and proliferation without inducing cell death. (A) Schematic overview of ex vivo proliferation and cell death analyses using *Nras^G12D^Ezh2^−/−^*BM. (B-C) Ex vivo proliferation of CD11b^+^ myelomonocytic cells from MEK inhibitor (MEKi)-treated BM (Mirdametinib or Trametinib; 1-100 nM, 48h). (B) Percentage of cells in the S phase following treatment with increasing doses of the MEKi Mirdametinib and Trametinib. n=3, analyzed with two-way ANOVA using Sidak’s correction for multiple comparisons comparing MEKi to vehicle control. (C) Representative pseudocolor plots of EdU incorporation. (D-E) Cell viability assays of total CD45^+^ cells from MEKi treated BM using Annexin V/7-AAD staining (Mirdametinib or Trametinib 1-100 nM, 24h). (D) Percentage of viable cells (Annexin V/7-AAD negative) following treatment with increasing doses of the MEKi Mirdametinib and Trametinib. n=3, analyzed with two-way ANOVA using Sidak’s correction for multiple comparisons comparing MEKi to vehicle control. (E) Representative pseudocolor plots of Annexin V/7-AAD staining. (F) Schematic overview of the ex vivo differentiation assays. (G-H) Myelomonocytic differentiation following lineage depletion and culture in the presence of GM-CSF (10 ng/mL) with vehicle or Trametinib (20 nM). (G) Quantification of CD11b^+^Gr1^+^ myelomonocytic cells. n=4; analyzed with two-way ANOVA using Sidak’s correction for multiple comparisons comparing MEKi to vehicle control. (H) Representative pseudocolor plots at baseline (Day 0) and after 7 days of treatment. (I) Schematic overview of in vivo MEKi treatment design. Mice were sacrificed after 10 weeks of treatment or when moribund. (J-K) Proliferation analysis by PCNA staining of spleens of *Nras^G12D^Ezh2^−/−^*mice sacrificed at the experimental endpoint. (J) Representative images of PCNA-stained spleen sections from vehicle-(left) and Mirdametinib-treated (right) mice. Scale bar represents 100 µM. (K) Quantification of PCNA^+^ cells. n=3; analyzed with unpaired *t* test. (L) Representative gating strategy for BM HSPC compartments. (M-N) Quantification of CMP (CD16/32^-^CD34^+^ LK cells), GMP (CD16/32^+^CD34^+^ LK cells) and MEP (CD16/32^-^CD34^-^ LK cells) cells in the BM of mice treated with Mirdametinib and Trametinib and sacrificed at the experimental endpoint. n= 5-11; analyzed with pairwise *t* test using correction for multiple testing. pIpC, polyinosinic:polycytidylic acid; MEKi, MEK inhibitor; PCNA, proliferating cell nuclear antigen; ns, not significant: CMP, common myeloid progenitor; GMP, granulocyte-monocyte progenitor; MEP, megakaryocyte-erythroid progenitor. Data are shown as mean ± SD; *p < .05, **p < .01, ***p < .001.

These findings were also corroborated in vivo (Figure 5I), where immunohistochemical staining of proliferating cell nuclear antigen (PCNA) demonstrated reduced proliferation in spleens of Mirdametinib-treated mice (Figure 5J-K). In agreement with our ex vivo differentiation data, flow cytometric analysis of the HSPC compartment in lineage-depleted BM revealed a significant reduction of GMPs in both Mirdametinib– and Trametinib-treated cohorts compared to vehicle-treated mice (Figure 5L-N).

### MEKi inhibits myeloid proliferation in primary *RAS^mut^EZH2^inac^*^t^ CMML patient specimens

To validate MAPK/ERK hyperactivation and MEKi sensitivity in human disease, we analyzed primary CMML patient specimens (Supplemental Table 1). We first performed immunohistochemical staining of phospho-MEK (p-MEK) and detected strong MAPK/ERK pathway activation in all *RAS^mut^EZH2^inact^* (3/3), compared to only a subset of *RAS^mut^* (2/4) and none of the *RAS^wt^* samples (0/5; Figure 6A). p-MEK signals were quantified according to a previously published scoring system.^46, 47^

**Figure 6:**
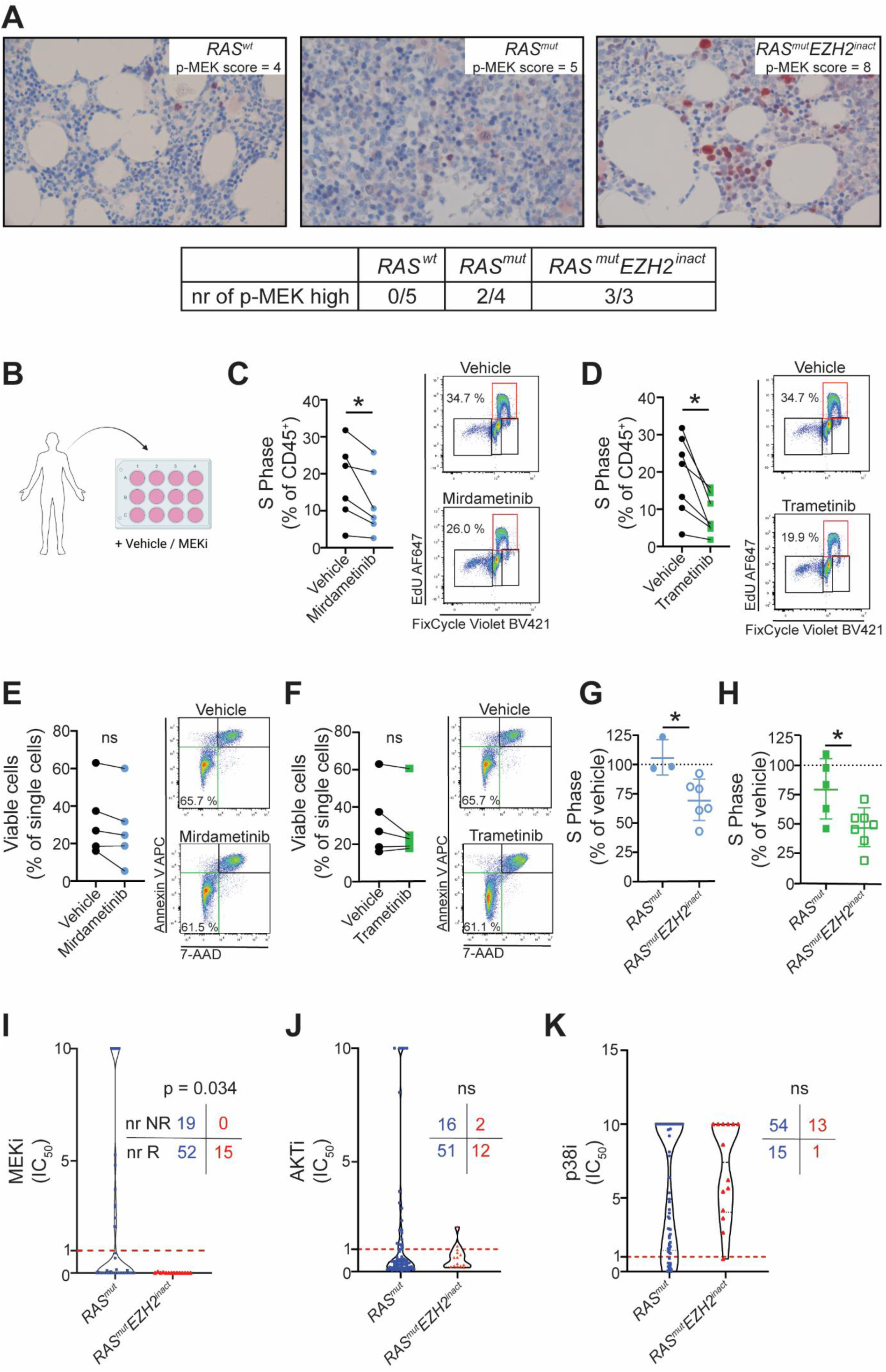
MEK inhibition suppresses myeloid proliferation in primary *RAS^mut^EZH2^inact^* CMML patient specimens. (A) Immunohistochemical analysis of p-MEK in primary patient samples with different genetic backgrounds. (B) Schematic overview of ex vivo analyses using primary patient-derived samples. (C-D) Ex vivo proliferation of MEK inhibitor (MEKi)-treated *RAS^mut^EZH2^inact^* primary patient specimens assessed by EdU incorporation. (C) Samples were treated with 20nM Mirdametinib for 48h. The percentage of CD45^+^ cells in S Phase following vehicle or Mirdametinib treatment and representative pseudocolor plots are shown. (D) Samples were treated with 20nM Trametinib for 48h. The percentage of CD45^+^ cells in S Phase following vehicle or Mirdametinib treatment and representative pseudocolor plots are shown. n=6-7; analyzed by Wilcoxon matched-pairs signed-rank test. (E-F) Ex vivo cell viability assays of MEKi treated *RAS^mut^EZH2^inac^*^t^ primary patient samples performed with Annexin V/7-AAD staining. (E) Samples were treated with 20nM Mirdametinib for 24h. The percentage of viable cells (Annexin V/7-AAD negative) and representative pseudocolor plots are shown. (F) Samples were treated with 20nM Trametinib for 24h. The percentage of viable cells (Annexin V/7-AAD negative) and representative pseudocolor plots are shown. n=5; analyzed by Wilcoxon matched-pairs signed-rank test. (G-H) Comparison of the anti-proliferative effects of Mirdametinib (G) and Trametinib (H) between *RAS^mut^* and *RAS^mut^EZH2^inact^* samples, shown as normalized decrease in proliferation. n=6-7; Data are shown as mean ± SD, analyzed with unpaired *t* test. (I-K) Analysis of ex vivo drug sensitivity in *RAS^mut^* and *RAS^mut^EZH2^inact^* AML patient specimens from the Beat-AML dataset.^37^ Samples were classified as responders (R; IC50≤1nM) or non-responders (NR; IC50>1nM). The analysis was performed for the MEKi Trametinib (I), the Akt inhibitor A674563 (J), and the p38α/MAPK14 inhibitor VX-745 (K). Responder frequencies between *RAS^mut^*and *RAS^mut^EZH2^inact^* groups were compared using a Fisheŕs exact test. ns, not significant.; * p < .05.

Ex vivo EdU-based proliferation assays (Figure 6B) demonstrated that both Mirdametinib (Figure 6C) and Trametinib (Figure 6D) significantly inhibited proliferation in *RAS^mut^EZH2^inact^*samples. Consistent with our findings in the mouse model, MEKi at the tested concentrations (20 nM, 24h) did not significantly induce cell death in primary patient specimens (Figure 6 E-F).

Direct comparison with *RAS^mut^* patient specimens revealed a significantly stronger anti-proliferative effect of MEKi in *RAS^mut^EZH2^inact^*specimens (Figure 6G-H), further confirming the genotype-specific sensitivity. Of note, our CMML cohort included samples obtained at diagnosis, relapsed/refractory disease, and even at secondary AML (sAML) progression (Supplemental Table 1). As sAML represents the most aggressive late-stage manifestation of CMML, we sought to determine whether this therapeutic vulnerability is translatable to sAML and AML in general. Therefore, we re-analyzed a previously published AML dataset comprising information about *RAS^mut^*and *EZH2^inact^* status, as well as ex vivo drug sensitivity data (n=86).^37^

Again, Trametinib exhibited significantly increased potency in *RAS^mut^EZH2^inact^* AML samples compared to *RAS^mut^*samples overall, with all *RAS^mut^EZH2^inact^* samples (15/15) showing IC_50_ values below 1nM, compared to 73% (52/71) of *RAS^mut^* specimens (Figure 6I). In contrast, inhibition of other RAS-downstream pathways, including PI3K/AKT (A674563) and p38/MAPK12 (VX-745), had no comparable effects (Figure 6J-K). This selective sensitivity to RAS-MAPK/ERK pathway inhibition in *RAS^mut^EZH2^inact^*genotypes is consistent with the preferential activation of this pathway in *Ras^mut^Ezh2^inact^* mice (Figure 2H and Supplemental Figure 2). To further support these observations, we re-analyzed the AML cohort for gene expression and assessed transcriptional signatures indicative of MAPK/ERK, PI3K/AKT or p38/MAPK activation. In more detail, pathway activation scores were calculated for each sample as median z-score of genes included in the BIOCARTA_ERK_PATHWAY, BIOCARTA_AKT_PATHWAY, and BIOCARTA_p38/MAPK_PATHWAY provided in the Molecular Signatures Database,^38, 39^ and then compared between *RAS^mut^EZH2^inact^*, *RAS^mut^*, and *RAS^wt^* cases. In these analyses, signatures indicative of MAPK/ERK activation increased in *RAS^mut^EZH2^inact^* samples, whereas gene signatures indicative of PI3K/AKT or p38/MAPK activation remained unchanged across respective genotypes (Supplemental Figure 3).

### MEKi suppresses expansion and proliferation of human *RAS^mut^EZH2^inact^* sAML transformed from CMML in a patient-derived xenotransplantation (PDX) model

To delineate the effects of MEKi on human *RAS^mut^EZH2^inact^* leukemic cells in vivo, we tested Trametinib in a PDX model established from a sAML sample transformed from CMML harboring an NRAS p.G13D mutation and deletion of chromosome 7. Following verification of engraftment, mice were randomized to receive Trametinib or vehicle control for 4 weeks (Figure 7A). Serial monitoring revealed significantly lower frequencies of human CD45^+^ cells (hCD45^+^) in the PB of Trametinib-treated mice compared to controls (Figure 7B–C). At the experimental endpoint, Trametinib treatment resulted in a reduced percentage of hCD45^+^ cells in spleen (Figure 7D) and total BM (Figure 7E), as well as a reduced percentage of hCD34^+^ cells in PB (Figure 7F). Of note, more than 99% of hCD45^+^ cells also expressed hCD33, confirming the myeloid origin of the engrafted leukemia (Supplemental Figure 4).

**Figure 7:**
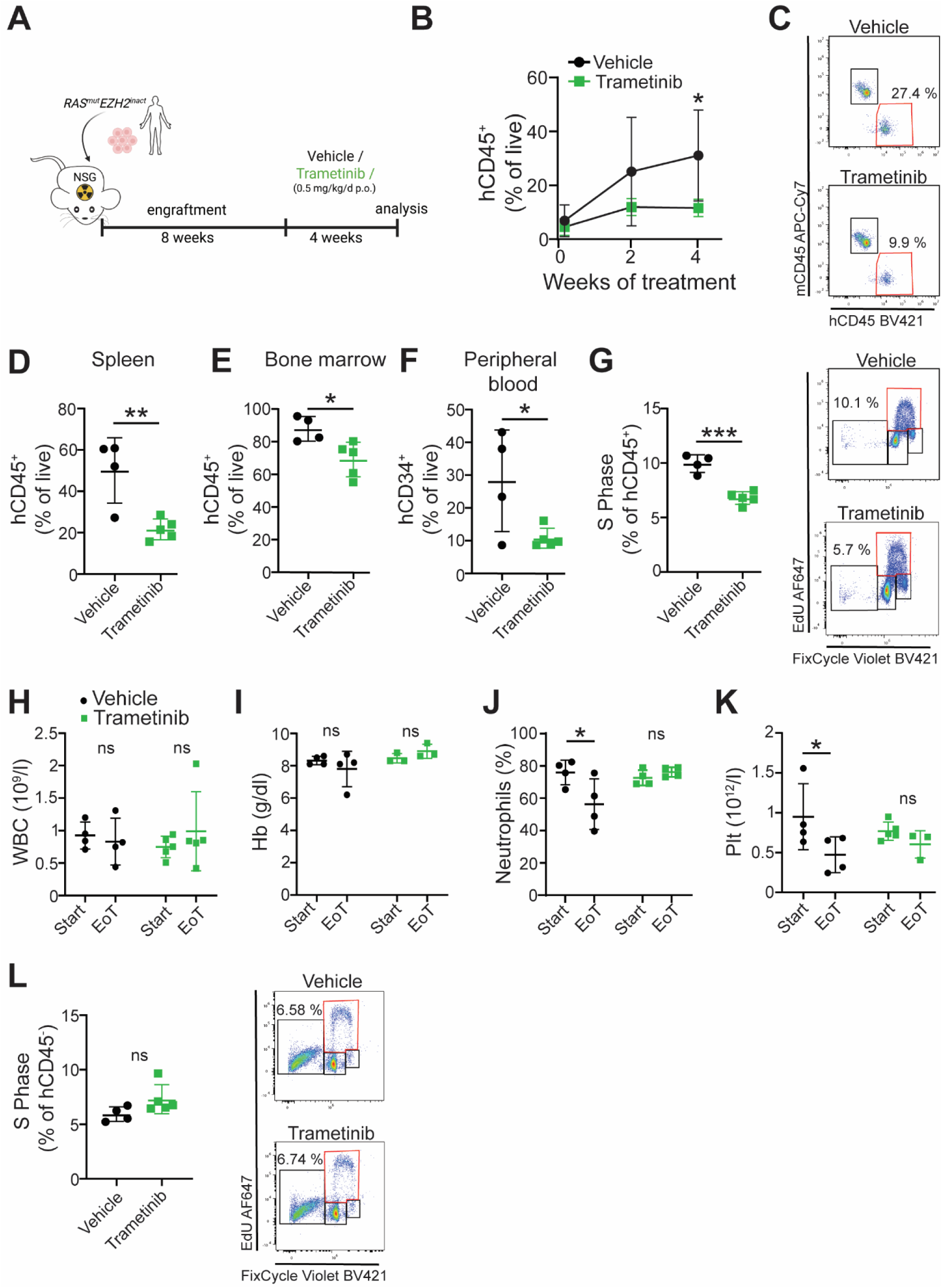
MEK inhibition suppresses expansion and proliferation of human *RAS^mut^EZH2^inact^* sAML transformed from CMML in a patient-derived xenotransplantation (PDX) model. (A) Schematic overview of the PDX treatment design. (B) Flow cytometric analysis of human (h)CD45^+^ cell engraftment in peripheral blood during treatment. n=4-5; analyzed by two-way ANOVA using Sidak’s correction for multiple comparisons. (C) Representative pseudocolor plots of mouse (m)CD45^+^ and hCD45^+^ cells in vehicle– (top) and Trametinib-treated (bottom) animals. (D-F) Leukemic burden at the end of the 4-week treatment period, as assessed by the percentage of hCD45^+^ in spleen (D) and total BM (E), and the percentage of hCD34^+^ cells in peripheral blood (F). Of note, more than 99% of hCD45^+^ cells also expressed hCD33, confirming the myeloid origin of the engrafted leukemia. (G) Ex vivo EdU-based proliferation assay of BM cells derived from vehicle– and Trametinib-treated mice at the experimental endpoint. Quantification of hCD45^+^ cells in S phase and representative corresponding pseudocolor plots are shown. n=4-5; analyzed with unpaired *t* test. (H-K) Peripheral blood counts at treatment start and end, including white blood cells (WBC, H) hemoglobin (Hb, I), neutrophils (J) and platelets (Plt, K). n=3-5; analyzed with two-way ANOVA using Sidak’s correction for multiple comparisons. (L) Ex vivo EdU-based proliferation assay of hCD45^−^ (non-leukemic) BM cells. Quantification of cells in S phase and representative pseudocolor plots are shown. n=4-5, analyzed with unpaired Student’s *t*-test. ns, not significant; EoT, end of treatment. Data are shown as mean ± SD;*p < .05, **p < .01, ***p < .001.

Ex vivo EdU-based proliferation assays performed on BM cells demonstrated a significant reduction of proliferating hCD45^+^ leukemic cells in Trametinib-treated mice (Figure 7G). This reduction of the transplanted leukemic clone in Trametinib-treated mice was further reflected in serial PB counts. Vehicle-treated mice developed progressive cytopenias, likely due to leukemic expansion and suppression of normal hematopoiesis, whereas these abnormalities were corrected in Trametinib-treated animals (Figure 7H–K).

Consistent with these findings, Trametinib selectively targeted *RAS^mut^EZH2^inact^* leukemic cells, as the proliferation of hCD45^-^ non-leukemic murine cells remained unaffected (Figure 7L). Moreover, no treatment-related toxicity was observed, as evidenced by stable body weight (Supplemental Figure 5A) and unchanged spleen morphology (Supplemental Figure 5B-D) in Trametinib-treated mice.

## Discussion

CMML is a heterogeneous hematologic malignancy with poor prognosis, particularly in patients ineligible for allogeneic transplantation.^4, 5^ Despite advances in molecular characterization, effective targeted therapeutic strategies remain limited. *RAS* mutations are among the most frequent genetic aberrations in CMML and are strongly associated with a proliferative disease phenotype; however, attempts to therapeutically target RAS-driven signaling have shown limited efficacy so far. We have previously demonstrated co-occurrence of *RAS^mut^* and *EZH2^inact^* within the Austrian Biodatabase for CMML.^28^ Moreover, we observed enhanced MAPK/ERK signaling and increased sensitivity to MEK inhibition in the *RAS^mut^EZH2^inact^*genotype in vitro.^28^ Building on these findings, we now establish the clinical and translational relevance of this genotype. By integrating patient cohorts and experimental models, we identify *RAS^mut^EZH2^inact^* as a high-risk subgroup characterized by aggressive disease biology, selective MAPK/ERK dependency, and pronounced sensitivity to MEK inhibition.

Initially, we re-analyzed the CMML subset of the IPSS-M cohort,^34^ and observed that *RAS^mut^EZH2^inact^*patients exhibit significantly shorter overall survival compared to those carrying *RAS^mut^* alone. These findings corroborate our previous observations within the Austrian Biodatabase for CMML.^28^ Similar to *RAS^mut^,*^12–14, 48, 49^ this genotype is enriched in MP-CMML and retains an adverse prognostic impact within this aggressive subtype. The aggressive nature of *RAS^mut^EZH2^inact^*was further recapitulated in a transgenic murine model, where *Nras^G12D^Ezh2^−/−^*results in a highly aggressive CMML-like myeloproliferative disease. These data are in agreement with previous data from Gu and coworkers^43^ and support the concept that *RAS* mutations in myeloid neoplasms require cooperating genetic alterations to fully manifest their leukemogenic potential. Indeed, co-occurrence of oncogenic *RAS* with loss of *Tet2*^27^ or *Rkip*^50–52^ has similarly been shown to enhance disease aggressiveness. In this context, *Ezh2* inactivation appears to act as a critical amplifier of oncogenic signaling, which is further supported by data showing disease aggravation in *Jak2^V617F^* myeloid neoplasms by additional *Ezh2* deletion.^53, 54^ Notably, *EZH2^inact^*in human patients can arise from both monoallelic and biallelic alterations,^40–42^ whereas our transgenic mouse model required complete loss of *Ezh2* to induce robust MAPK/ERK hyperactivation. This discrepancy likely reflects the more uniform and less complex (epi-)genetic background of murine models compared to human disease. While we addressed this limitation by focusing on mice with complete *Ezh2* deletion and by incorporating PDX models, these findings highlight the need for more complex genetically engineered models to more accurately reflect the human disease.

At the biological level, we demonstrate that co-occurrence of *RAS^mut^*and *EZH2^inact^* results in selective hyperactivation of MAPK/ERK signaling. This was consistently observed in all murine models and human patient specimens studied, and even held true in transcriptomic profiling of a large AML dataset,^37^ suggesting that enhanced MAPK/ERK signaling may represent a common downstream consequence of this mutational combination not only in CMML but in myeloid leukemogenesis in general. Importantly, MAPK/ERK activation was detectable not only in bulk tumor populations but also within the majority of stem and myeloid progenitor cell compartments. Considering previous data that activated MAPK/ERK signaling induces myelomonocytic differentiation^55^ and proliferation of undifferentiated hematopoietic cells,^56^ these data support a model in which *RAS^mut^EZH2^inact^* contributes to CMML development already at very early stages of hematopoiesis. In contrast, other RAS effector pathways, including PI3K/AKT and p38/MAPK, were not activated, indicating selective activation of MAPK/ERK signaling. These data are in contrast to TET2 loss in the context of oncogenic NRAS, where their co-occurrence induced the activation of both MAPK/ERK and PI3K/AKT signaling,^27^ highlighting how combinations of different epigenetic and signaling mutations can lead to context-dependent differences in downstream signaling.

This selective MAPK/ERK dependency provides a strong rationale for therapeutic targeting. Therefore, we evaluated MEKi in the *Nras^G12D^Ezh2^inact^*^-^driven murine leukemia model and treated the mice with the orally available MEKi Mirdametinib and Trametinib. These drugs were chosen as they selectively target MAPK/ERK signaling and due to their broad clinical use, making them a directly translatable therapeutic option for affected patients. MEKi effectively suppressed MAPK/ERK activation in both mature myeloid cells as well as hematopoietic stem and myeloid progenitor compartments, indicating the reversal of the aberrant *RAS^mut^EZH2^inact^*-induced signaling phenotype at all stages of hematopoietic development. Consequently, MEKi induced a profound anti-leukemic effect, as evidenced by prolonged survival and reduction of myeloproliferation and leukocytosis. Although the *Nras^G12D^Ezh2^inact^* mice exhibit Mx1-Cre driven continuous expression of the mutant genotype in all hematopoietic cells, MEKi led to sustained stabilization of disease parameters, further supporting a dependency on MAPK/ERK signaling in this setting. The anti-leukemic activity of MEK inhibition was further validated in ex vivo-treated primary *RAS^mut^EZH2^inact^* CMML patient samples, including samples obtained at diagnosis, as well as during disease progression, treatment resistance, and secondary AML transformation. These advanced disease stages are typically associated with poor prognosis and limited therapeutic options, underscoring the potential clinical relevance of this approach. Consistently, analysis of the ex vivo drug sensitivity data within the Beat-AML cohort demonstrated increased sensitivity of *RAS^mut^EZH2^inact^*samples to MEKi compared to *RAS^mut^* samples alone.

Mechanistically, MEKi predominantly suppressed proliferation and myelomonocytic differentiation, while not affecting cell death. These data are in agreement with the established role of MAPK/ERK signaling in promoting mitogenic signaling and cell cycle progression.^57^ At a transcriptional level, MEKi was associated with suppression of oncogenic signaling programs, but also with upregulation of metabolic pathways, including oxidative phosphorylation, and MYC-associated transcriptional programs. This is particularly relevant as MEKi-treated *RAS^mut^* cancer cells have been shown to reduce glycolysis and activate oxidative phosphorylation as a resistance mechanism to evade apoptosis.^58, 59^ Similarly, upregulation of a MYC-dependent transcriptional program resulting in apoptosis evasion was reported in drug resistance following MEKi.^60^ Therefore, MEKi use may expose a metabolic shift and vulnerability that can be targeted with anti-apoptotic drugs, potentially improving long-term therapeutic outcomes. Although these data are preliminary, they highlight potential mechanisms of MEKi resistance in *RAS^mut^EZH2^inact^* CMML, thereby providing potential avenues to explore in future studies.

An important observation arose from the PDX model employed, where MEKi suppressed proliferation of the *RAS^mut^EZH2^inact^* engrafted human leukemia cells, while proliferation within residual non-malignant murine hematopoiesis was unchanged. These data suggest that MEKi selectively targets *RAS^mut^EZH2^inact^*leukemic cells while sparing non-malignant hematopoiesis in vivo, indicating a potential therapeutic window. This hypothesis is further supported by the above-mentioned data from the ex vivo treatments, where *RAS^mut^EZH2^inact^* cases exhibited a preferential sensitivity to MEKi when compared *RAS^mut^*alone. These data are of particular relevance for the treatment of human CMML patients with MEKi. Although these drugs are already available for the clinical routine setting, the enthusiasm for their use in *RAS^mut^* CMML have previously been dampened by their modest success in clinical trials of adult *RAS^mut^* leukemia.^21, 61^ Although some patients demonstrated good responses, the vast majority did not benefit sufficiently from this therapy. Our findings now provide a mechanistic explanation for the failure of MEKi in unselected CMML. They strongly suggest that these studies did not account for co-occurring genetic and epigenetic alterations and therefore likely included patients without MAPK/ERK pathway dependency. Our data suggest that molecular stratification – specifically the identification of *RAS^mut^EZH2^inact^*cases — may be essential to unlock the clinical potential of MEKi in CMML. Another interesting approach for this aggressive patient subgroup arises from the recent development of direct RAS inhibitors. While KRAS-specific inhibitors already entered clinical routine,^62^ broader panRAS inhibitors were recently developed as well.^63, 64^ However, considering their early stage of clinical development, they do not represent an option for direct clinical use. Moreover, RAS(ON) inhibitors affect multiple downstream pathways, whereas our data demonstrate that *RAS^mut^EZH2^inact^*leukemias exhibit a selective dependency on MAPK/ERK signaling. In this context, targeted MEKi may provide a more specific approach, potentially even reducing unwanted side effects. Future studies will be necessary to validate this assumption. They might even clarify whether a combination of both therapeutic approaches further increases efficacy and/or opens a window for overcoming therapeutic resistance.

In summary, our study defines *RAS^mut^EZH2^inact^* CMML as a biologically and clinically distinct subgroup characterized by an aggressive disease course and selective hyperactivation of RAS – MAPK/ERK signaling. Using mouse, patient, and PDX models, we demonstrate that this genotype exhibits pronounced sensitivity to MEKi, primarily through suppression of proliferative and myelomonocytic expansion programs. Most importantly, these findings define a strong rationale for genotype-guided clinical evaluation of MEKi in *RAS^mut^*CMML, incorporating not only the *RAS^mut^* status, but also the presence of *RAS^mut^* co-occurring genetic and epigenetic events.

## Acknowledgements

This project was supported by Biobank Graz of the Medical University of Graz, Austria (Cohort 6002_14, Myeloid Neoplasm Collection). AZ is funded by the Austrian Science Fund (FWF, Grant-DOI 10.55776/P36672 and Grant-DOI 10.55776/PAT1753824), and the ERA-NET TRANSCAN-3 initiative (Austrian Science Funds Grant-DOI 10.55776/I6101). Research in his laboratory and Leukemia biobanking at the Medical University of Graz is further supported by Leukämiehilfe Steiermark. Students PC and AK are funded by the FWF within the PhD program Molecular Medicine of the Medical University of Graz. Student MCM is funded by the PhD program Molecular Medicine of the Medical University of Graz. AR was supported by grants from the Austrian Science Fund (FWF, Grant-DOI 10.55776/P32783 and Grant-DOI 10.55776/I5021) and received support from the Austrian Society of Hematology and Oncology (Clinical Research Grant), Leukämiehilfe Steiermark and MEFOgraz. The Figures were in part created with Biorender (https://biorender.com). ChatGPT (OpenAI; https://chatgpt.com/) and Grammarly (https://www.grammarly.com/) were used to improve the clarity and grammar of the manuscript. All outputs were carefully reviewed, revised, and integrated by the authors, who take full responsibility for the final content.

## Authorship

Contribution: AZ and EG designed and supervised the study; EG, BP, AK, PC, KL, MCM, JF, MK, GH and KK performed the research; SD, SW, JN, KG, AH, AW, HS, AR and AZ contributed patient specimens, vital material and/or collected clinical data; EG, BP, AK, PC, KL, MCM, JF, MK, KVC, GH, GB, GP and AZ collected data and performed statistical analyses; EG, BP, AK, PC, KL, MCM, JF, MK, KVC, KK, KG, AH, AW, HS, AR and AZ analyzed and interpreted the data; EG and AZ wrote the manuscript; All authors read, revised and approved the manuscript.

