## Supplemental Information for "EZH2 inactivation drives MAPK-dependent vulnerability to MEK inhibition in RAS-mutant CMML"

**Gruden E., et al.**

### Supplemental table 1: Patient characteristics

| Patient ID | Sex | Sample ID | Disease Stage at sampling | Age at Sample (y) | Prior therapy | Sample Source | Karyotype | Mutation status | Group | Ex vivo testing | pMEK BM Immunohostochemistry |
|---|---|---|---|---|---|---|---|---|---|---|---|
| LB-MUG-137 | f | 7050 and 7378/2011 | CMML PD or r/r | 68 | Hydroxyurea | PB and BM | 45,XX,-7 | CBL p.G413D, VAF 97%<br>EZH2 p.Q553del, VAF 89% | $RAS^{mut}EZH^{inact}$ | yes | yes |
| LB-MUG-289 | f | 8686 | sAML from CMML | 58 | no | BM | 46,XX,t(6;11)(q27;q23), del(7q) | IDH2 p.R140Q, VAF 11%<br>KRAS p.G13C, VAF 30%<br>NRAS p.G13D, VAF 5% | $RAS^{mut}EZH^{inact}$ | yes | no material |
| LB-MUG-294 | m | 9402 | sAML from CMML | 56 | no | PB | 45,XY,inv(3)(q21q26),-7 | BCOR:exon7:c.3239-2A>G:splice-site-acceptor-mutation, VAF 72%<br>CEBPA p.P197delinsHPP, VAF 41%<br>FLT3-ITD ratio of 0,12<br>NRAS p.G13D, VAF 48%<br>SRP72 p.G7E, VAF 48%<br>U2AF1 p.S34F, VAF 42% | $RAS^{mut}EZH^{inact}$ | yes | no material |
| LB-MUG-355 | m | 10183 and 3050/2020 | CMML PD or r/r | 72 | Hydroxyurea | PB and BM | 44,XY,-Y,-7 | CBL p.L380P, VAF 31%<br>DNMT3A p.P799S, VAF 39%<br>DNMT3A p.I695S, VAF 37%<br>IDH2 p.R140Q, VAF 39%<br>JAK2 p.V617F, VAF 18%<br>NF1 p.Q1617*, VAF 3%<br>TET2 p.P174H, VAF 51% | $RAS^{mut}EZH^{inact}$ | yes | yes |
| LB-MUG-358 | m | 8843 | CMML Dg | 74 | no | PB | 46,XY | EZH2 p.L674F, VAF 98 %<br>NRAS p.G12D, VAF 46%<br>RUNX1 p.T312fs, VAF 47%<br>TET2 p.C1135Y, VAF 48%<br>TET2 p.T1372I, VAF 47% | $RAS^{mut}EZH^{inact}$ | yes | no material |
| LB-MUG-573 | f | 9467 and 4305/2019 | CMML Dg | 58 | no | PB and BM | 45,XX,-7 | CBL p.I383M, VAF 31%<br>DNMT3A p.F731fs, VAF 45%<br>IDH2 p.R140Q, VAF 47%<br>NRAS p.G13D, VAF 13%<br>NRAS p.G12V, VAF 20% | $RAS^{mut}EZH^{inact}$ | yes | yes |
| LB-MUG-573 | f | 9591 | sAML from CMML | 58 | 7+3 CTX | PB | 47,XX,+4,+11,-7 | CBL p.I383M, VAF 31%<br>DNMT3A p.F731fs, VAF 45%<br>IDH2 p.R140Q, VAF 47%<br>NRAS p.G13D, VAF 13%<br>NRAS p.G12V, VAF 20% | $RAS^{mut}EZH^{inact}$ | yes | no material |
| LB-MUG-115 | m | 9465 and 36011/2019 | CMML Dg | 58 | no | PB and BM | 46,XY | PTPN11 p.A72G, VAF 3% | $RAS^{mut}$ | yes | yes |
| LB-MUG-138 | m | 8849 | CMML Dg | 81 | no | BM | 47,XY,+8 | ASXL1 p.D741fs, VAF 34%<br>DNMT3A p.T437M, VAF 51%<br>NRAS p.G12D, VAF 46%<br>SRSF2 p.P95H, VAF 44%<br>TET2 p.C1289*, VAF 94% | $RAS^{mut}$ | yes | no material |

Supplemental table 1: Patient characteristics… continued from previous page

| Patient ID | Sex | Sample ID | Disease Stage at sampling | Age at Sample (y) | Prior therapy | Sample Source | Karyotype | Mutation status | Group | Ex vivo testing | pMEK BM Immunohostochemistry |
|---|---|---|---|---|---|---|---|---|---|---|---|
| LB-MUG-231 | f | 9284 | CMML Dg | 69 | no | BM | 46,XX | CBL p.C381F, VAF 29%<br>DNMT3A p.R882H, VAF 44%<br>RUNX1 p.R204*, VAF 3% | $RAS^{mut}$ | yes | no material |
| LB-MUG-336 | m | 73047/2018 | CMML Dg | 69 | no | BM | 46,XY | ASXL1 p.W583*, VAF 45%<br>CBL p.C384Y, VAF 33%<br>IDH2 p.R140Q, VAF 48%<br>SRSF2 p.P95H, VAF 45% | $RAS^{mut}$ | no materi | yes |
| LB-MUG-365 | m | 74416/2017 | CMML Dg | 42 | no | BM | 46,XY | KRAS p.G12D, VAF 39% | $RAS^{mut}$ | no materi | yes |
| LB-MUG-427 | f | 36253/2022 | CMML Dg | 63 | no | BM | 46,XX | KRAS p.G12D, VAF 38%<br>ZRSR2 p.R451_S455dup, VAF 33% | $RAS^{mut}$ | no materi | yes |
| LB-MUG-464 | m | 9315 | CMML Dg | 78 | no | BM | 46,XY | ASXL1 p.S833fs, VAF 37%<br>CBL p.Y371H, VAF 84%<br>PHF6 p.R342*, VAF 84%<br>TET2 p.P656fs, VAF 42%<br>TET2 p.E1483*, VAF 44%<br>ZRSR2 exon11:splice_site_acceptor_mutation,c.938G>A, VAF 87% | $RAS^{mut}$ | yes | no material |
| LB-MUG-712 | m | 10763 | CMML PD or r/ | 74 | Azacitidine | PB | 47,XY,+8 | CBL p.D390H, VAF 6%<br>IDH2 p.R140Q, VAF 43%<br>NRAS p.G12V, VAF 41%<br>SRSF2 p.P95R, VAF 43% | $RAS^{mut}$ | yes | no material |
| LB-MUG-119 | m | 86846/2014 | CMML Dg | 73 | no | BM | 46,XY | JAK2 p.V617F, VAF 83%<br>TET2 p.Q531X, VAF 46%<br>TET2 p.M611fs, VAF 51% | $RAS^{wt}$ | no materi | yes |
| LB-MUG-149 | m | 103661/2016 | CMML Dg | 74 | no | BM | 46,XY | U2AF1 p.Q157P, VAF 48% | $RAS^{wt}$ | no materi | yes |
| LB-MUG-157 | m | 3807/2021 | CMML Dg | 55 | no | BM | 46,XY | SRSF2 p.P95H, VAF 45%<br>TET2 p.Q531*, VAF 47% | $RAS^{wt}$ | no materi | yes |
| LB-MUG-360 | f | 36942/2014 | CMML Dg | 45 | no | BM | 46,XX | IDH1 p.R132C, VAF 49%<br>SF3B2 p.A26S, VAF 48% | $RAS^{wt}$ | no materi | yes |
| LB-MUG-541 | m | 50994/2014 | CMML Dg | 66 | no | BM | 46,XY | TET2 p.K1090fs, VAF 78% | $RAS^{wt}$ | no materi | yes |

PANEL 1 (Supplemental Table 2)

| **Antibody** | **Dilution** | **Clone** | **Company** | **Cat#** | **RRID** |
|---|---|---|---|---|---|
| CD11b efluor450 | 1:150 | M1/70 | eBioscience | 48-0112-82 | AB_1582236 |
| Ly6G/Ly6C PE-Cy7 | 1:150 | RB6-8C5 | eBioscience | 25-5931-82 | AB_469663 |
| CD115 PE | 1:150 | AFS98 | eBioscience | 12-1152-82 | AB_465808 |
| CD117 APC | 1:50 | 2B8 | BD Pharmingen | 553356 | AB_398536 |
| CD45 FITC | 1:75 | 30-F11 | eBioscience | 11-0451-82 | AB_465050 |
| B220 APC-Cy7 | 1:150 | RA3-6B2 | BD Pharmingen | 552094 | AB_394335 |
| DX5 APC-Cy7 | 1:20 | DX5 | eBioscience | 47-5971-82 | AB_11218895 |
| CD3e APC-Cy7 | 1:20 | 145-2C11 | BD Pharmingen | 557596 | AB_396759 |

PANEL 2 (Supplemental Table 3)

| **Antibody** | **Dilution** | **Clone** | **Company** | **Catalogue #** | **RRID** |
|---|---|---|---|---|---|
| CD48 FITC | 1:37.5 | HM48-1 | Biolegend | 103403 | AB_313018 |
| CD150 BV421 | 1:10 | TC15-12F12.2 | Biolegend | 115925 | AB_10896787 |
| Sca1 PE-Cy7 | 1:20 | D7 | eBioscience | 25-5981-82 | AB_469669 |
| CD117 APC | 1:150 | 2B8 | BD Pharmingen | 553356 | AB_398536 |

PANEL 3 (Supplemental Table 4)

| **Antibody** | **Dilution** | **Clone** | **Company** | **Catalogue #** | **RRID** |
|---|---|---|---|---|---|
| CD117 APC | 1:150 | 2B8 | BD Pharmingen | 553356 | AB_398536 |
| Sca1 PE-Cy7 | 1:20 | D7 | eBioscience | 25-5981-82 | AB_469669 |
| CD16/32 eFluor 450 | 1:75 | 93 | eBioscience | 48-0161-82 | AB_1272191 |
| CD34 PE | 1.10 | RAM34 | BD Pharmingen | 551387 | AB_394176 |

PANEL 4 (Supplemental Table 5)

| Antibody | Dilution | Clone | Company | Catalogue # | RRID |
|---|---|---|---|---|---|
| anti-mouse CD45 APC-Cy7 | 1:50 | 30-F11 | BD Pharmingen | 557659 | AB_396774 |
| anti-mouse Ter119 PE-Cy5 | | TER-119 | Biolegend | 116210 | AB_313711 |
| anti-human CD45 BV421 | | HI30 | Biolegend | 304032 | AB_2561357 |
| anti-human CD33 PE | | WM53 | eBioscience | 12-0338-42 | AB_10855036 |
| anti-human CD19 PE-Dazzle594 | | SJ25C1 | Biolegend | 363032 | AB_2616853 |
| anti-human CD3 PE-Cy7 | | UCHT1 | Biolegend | 300420 | AB_439781 |
| anti-human CD34 FITC | | 581 | BD Pharmingen | 555821 | AB_396150 |

PANEL 5 (Supplemental Table 6)

| Antibody | Dilution | Clone | Company | Catalogue # | RRID |
|---|---|---|---|---|---|
| CD34 PE | 1:10 | RAM34 | BD Pharmingen | 551387 | AB_394176 |
| Sca-1 Pacific Blue | 1:40 | D7 | Biolegend | 108120 | AB_493273 |
| CD117 APC | 1:80 | 2B8 | BD Pharmingen | 553356 | AB_398536 |
| Fixable Viability Dye eFluor506 | 1:10 | | eBioscience | 65-0866-14 | |

PANEL 6 (Supplemental Table 7)

| **Antibody** | **Dilution** | **Clone** | **Company** | **Catalogue #** | **RRID** |
|---|---|---|---|---|---|
| CD45 PE-Cy7 | 1.20 | HI30 | Biolegend | 982310 | AB_2715773 |
| CD34 FITC | 1:10 | 581 | BD Pharmingen | 555821 | AB_396150 |
| CD38 PE | 1:20 | HIT2 | Biolegend | 980302 | AB_2616620 |

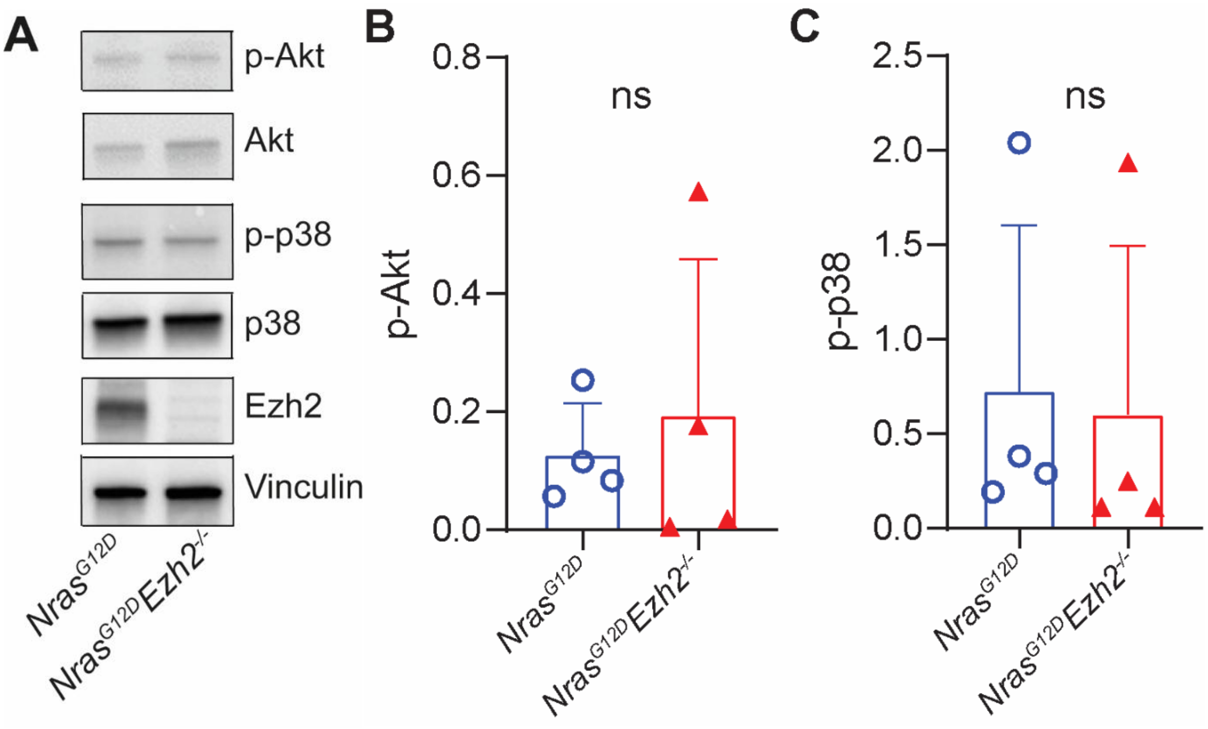


**Supplemental Figure 1: *Nras$^{G12D}$Ezh2$^{-/-}$* mice do not exhibit differences in PI3K/Akt or p38/MAPK pathway activation.** (A) Representative Immunoblot and quantification of (B) p-Akt and (C) p-p38 levels in *Nras$^{G12D}$* and *Nras$^{G12D}$Ezh2$^{-/-}$* mice. n=3-4; analyzed with unpaired Student´s *t*-test. ns, not-significant. Data is shown as mean + SD.

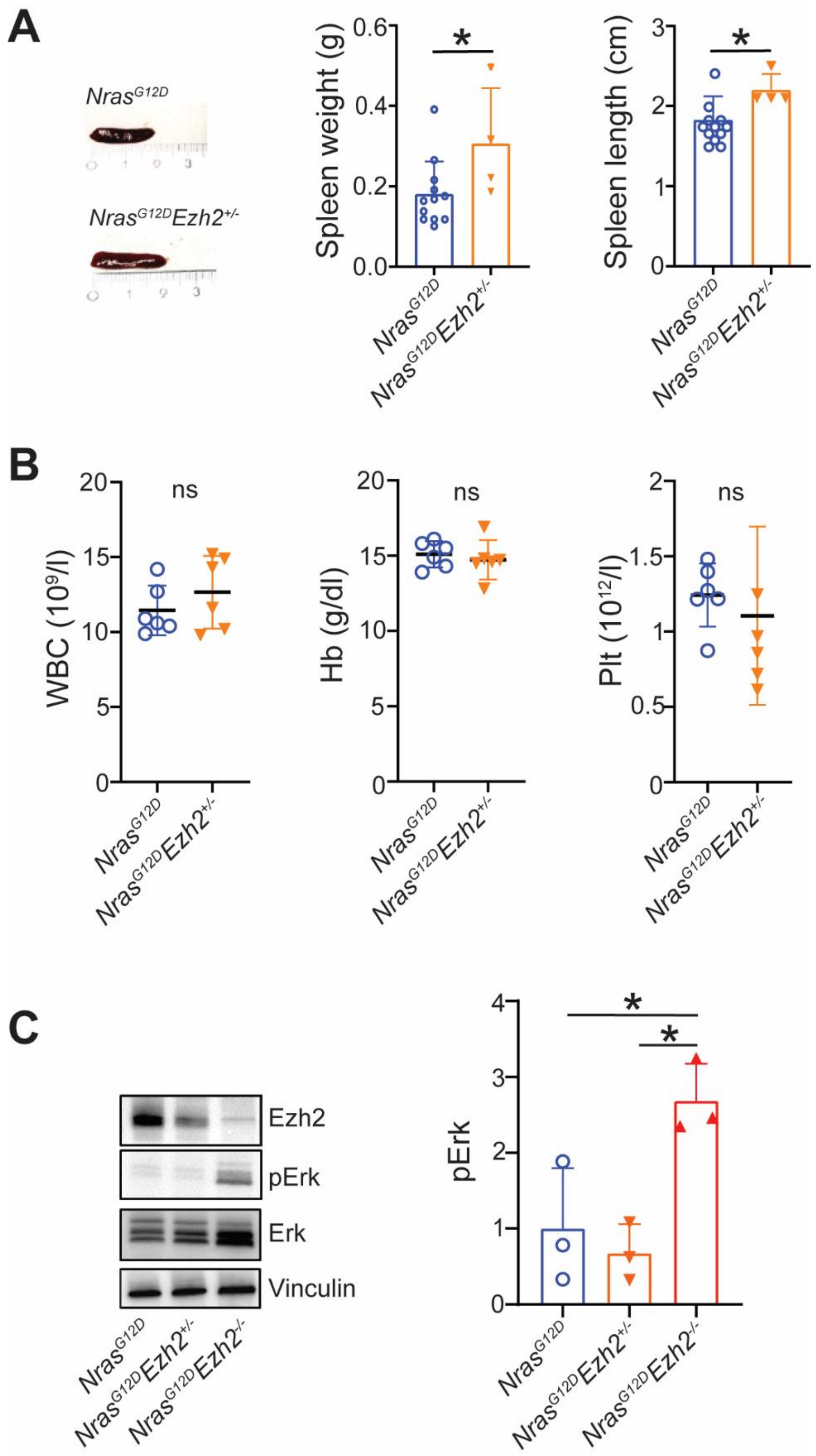


**Supplemental Figure 2: *Nras*$^{G12D}$*Ezh2*$^{+/-}$ mice do not develop an aggressive phenotype and lack overactivation of the MAPK/ERK pathway.** (A) Representative images and quantification of spleen weight and length of *Nras*$^{G12D}$ and *Nras*$^{G12D}$*Ezh2*$^{+/-}$ mice. Groups were compared by unpaired Student´s *t*-test. (B) Peripheral blood counts, including white blood cells (WBC), hemoglobin (Hb) and platelets (Plt), measured 15 weeks post-pIpC. n=4-12; analyzed with unpaired Student´s *t*-test. (C) Representative Immunoblot and quantification of pErk levels in total bone marrow from $Nras^{G12D}$, $Nras^{G12D}Ezh2^{+/-}$ and $Nras^{G12D}Ezh2^{-/-}$ mice. Comparisons were performed in three independent experiments using one-way ANOVA with Sidak´s multiple comparisons test.; ns, not significant. Data is shown as mean ± SD; *p < .05.

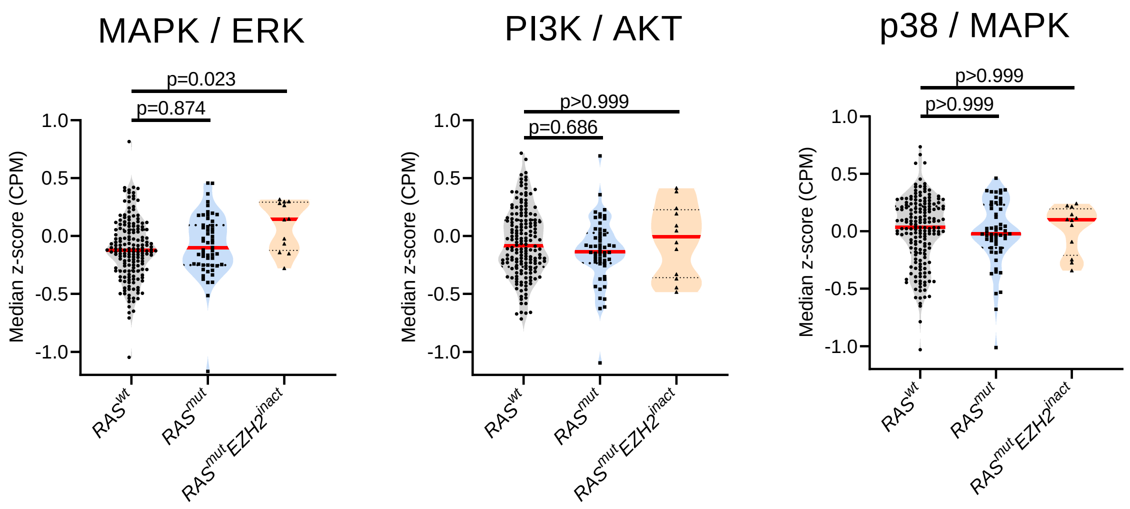


**Supplemental Figure 3: Selective upregulation of MAPK/ERK signaling in *$RAS^{mut}EZH2^{inact}$* patient samples.** The Beat-AML dataset[1], previously used for drug sensitivity analysis, was re-analyzed for transcriptional signatures indicative of MAPK/ERK, PI3K/AKT or p38/MAPK activation. In more detail, pathway activation scores were calculated for each sample as median z-score of genes included in the BIOCARTA_ERK_PATHWAY, BIOCARTA_AKT_PATHWAY and BIOCARTA_p38/MAPK_PATHWAY gene sets,[2, 3] and then compared between $RAS^{mut}EZH2^{inact}$, $RAS^{mut}$ and $RAS^{wt}$ cases. Data were analyzed using Kruskal–Wallis test followed by Dunn’s multiple comparisons test. Abbreviations: CPM, counts per million. Data are shown as violin plots with median and interquartile range.

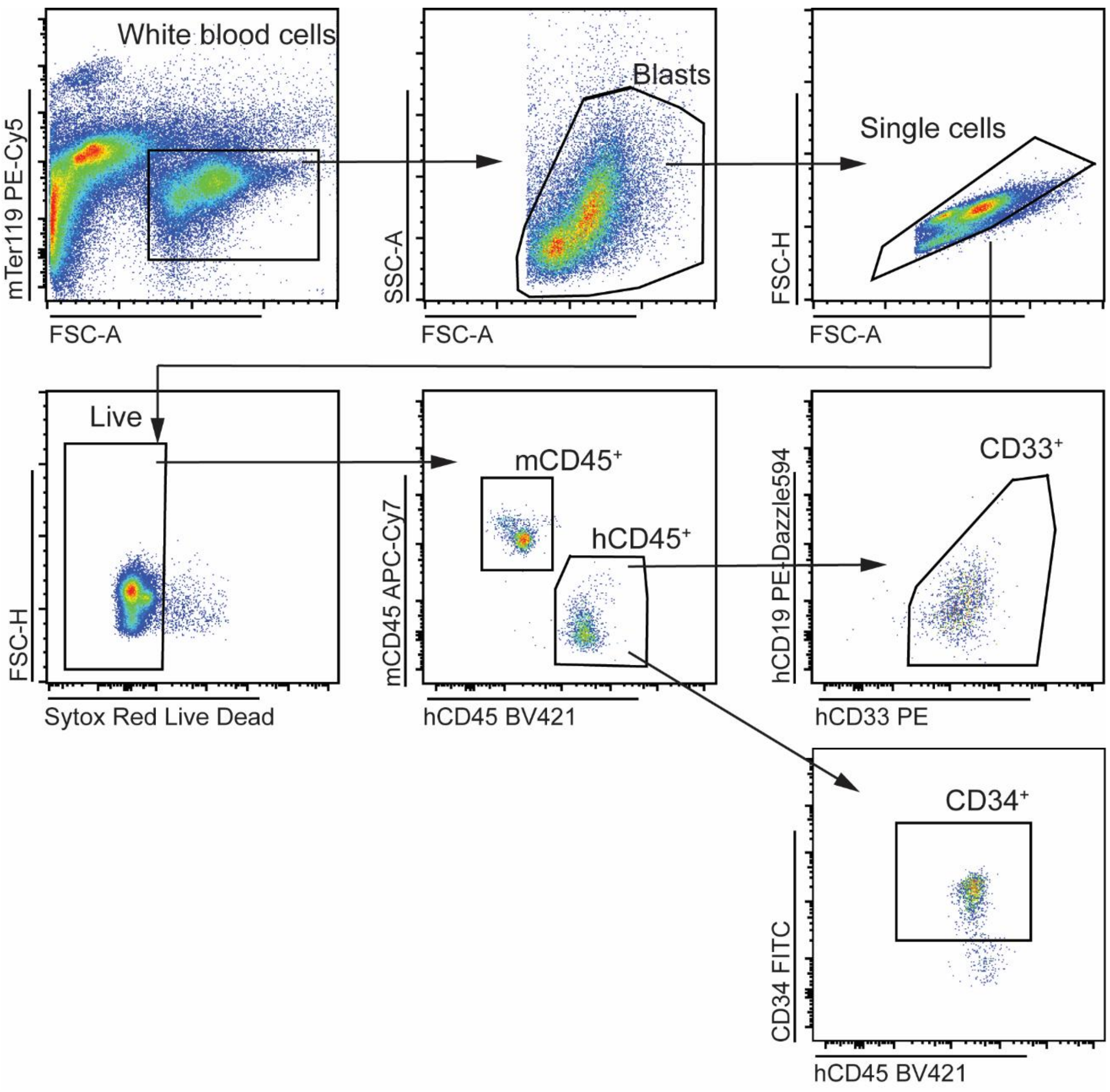


**Supplemental Figure 4: Gating strategy for PDX sample analysis.** Detailed flow cytometric staining panel for PDX sample analysis is found in Supplemental Table 5.

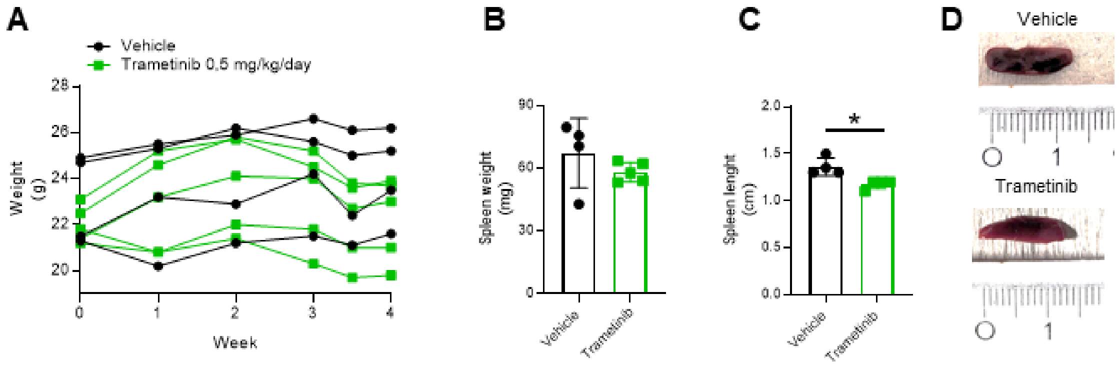


**Supplemental Figure 5: MEK inhibition does not induce toxicity in a PDX model of sAML.** (A) Body weight tracking of individual vehicle- or Trametinib-treated engrafted mice over the treatment period. At the end of the treatment period spleen weight (B) and length (C) were evaluated. n= 4-5; analyzed with unpaired Student´s *t*-test. (D) Representative images of spleens from vehicle- (top) and Trametinib-treated (bottom) mice. Data is shown as mean ± SD; *p < .05.

#### 1. Material and methods

##### 1.1 Transgenic Mice

###### 1.1.1 Routine blood counts

Blood from mice was collected via cheek bleeding into EDTA-coated tubes. Peripheral blood counts were performed on a Vet Hematology Analyzer (V–SIGHT; A. Menarini Pharma GmbH, Vienna, Austria) according to established protocols.

###### 1.1.2 Treatment of mice

10-12 week old Nras$^{G12D}$-Ezh2$^{-/-}$ mice were treated either with Mirdametinib (1.5 mg/kg/d per os)[4] or Trametinib (0.5 mg/kg/d per os)[5] or vehicle control (0.5% w/v Hydroxyl-propyl-methyl-cellulose + 0.1% Tween80 in deionized water) for 10 weeks. The long-term survival cohort was treated with Mirdametinib for 5 months (1.5 mg/kg/d per os). Survival and health status of treated mice was monitored daily. Mice were sacrificed when predefined clinical signs of sickness were observed (pain behavior and hemoglobin blood count values under 8 g/dl). Peripheral blood counts and flow cytometry were assessed at the start and bi-weekly for the duration of treatment or once monthly in the long term treatment cohort. At sacrifice, we measured spleen weight, length and peripheral blood counts. Lineage-depleted bone marrow, spleen and peripheral blood cells were additionally analyzed by flow cytometry. Histology and immunohistochemistry were performed from spleen samples. For phospho-flow characterization of mouse bone marrow progenitor compartments and total RNA sequencing analysis, mice were treated with Mirdametinib (1.5 mg/kg/d per os) or Trametinib (0.5 mg/kg/d per os) or vehicle control for 10 days.

###### 1.1.3 Single cell suspensions

Bone marrow cells from spine, pelvic bones, femur and tibia were collected by crushing with a mortar in $Ca^{2+}$ and $Mg^{2+}$-free Hank's buffered salt solution (HBSS; Gibco). Following cell lysis (BD PharmLyse, 5min, 4 °C), cells were strained through a 70 µM cell strainer. Spleens were dissociated via meshing through a 70 µM cell strainer. Cell numbers were determined with Casy cell counter (Schafer Systems, Model TTC).

##### 1.1.4 Ex vivo cell death and cell cycle analysis of $Nras^{G12D}$-$Ezh2^{-/-}$ bone marrow

Single cell suspensions of bone marrow were resuspended in IMDM supplemented with 10% FCS (FBS Good Forte, P40-47500-HI; PAN-Biotech), 1% Antibiotic-Antimycotic (100X, Thermo Scientific) and 10ng/ml of mGM-CSF in a 24 well plate at a cell density of $1x10^6$ per ml.

For cell death analysis cells were incubated with either MEKi (Mirdametinib or Trametinib, 1-100nM, 24h) or DMSO vehicle control. Following the incubation period, cells were washed and resuspended in Annexin binding buffer (BD Pharmingen) with the addition of Annexin V (1:20) and 7-AAD (1:20). Analysis of viable (Annexin $V^-$/7-$AAD^-$) cells was performed on a flow cytometer within 1h.

For cell cycle/proliferation analysis, cells were incubated with either MEKi (Mirdametinib or Trametinib, 1-100nM, 48h) or DMSO vehicle control. MEKi and DMSO vehicle were refreshed after 24h. Following the incubation period, each specimen was incubated with EdU (10µM, Click-iT™ Plus EdU Alexa Fluor™ 647 Flow Cytometry Assay Kit, Thermo Fisher Scientific, Waltham, MA, USA, #C10634). 1-2 hours later, cells were harvested and stained with anti-mouse CD11b PE (1:100; BD Pharmingen Cat #557397; 15min, RT). FxCycle Violet (1:1000 dilution, Thermo Fisher Scientific, #F10347) was added prior to acquisition with a flow cytometer.

##### 1.1.5 Lineage depletion using the Hematopoietic Progenitor Cell Enrichment Set

Hematopoetic progenitor cells from mouse bone marrow were enriched using the Mouse Hematopoetic Progenitor Cell enrichment set (BD, Bioscience, Cat.#558451) according to manufacturer's protocol and used for HSC/HSPC flow cytometric analysis or in ex vivo differentiation assays.

###### 1.1.6 Ex vivo differentiation assays

Lineage depleted *Nras*$^{G12D}$ *Ezh2*$^{-/-}$ bone marrow cells were resuspended in IMDM supplemented with 10% FCS (FBS Good Forte, P40-47500-HI; PAN-Biotech ), 1% Antibiotic-Antimycotic (100X, Thermo Scientific) and 10ng/ml of mouse (m)GM-CSF in a 24 well plate at a cell density of 0.3-0.6x$10^6$ per ml. MEKi (Trametinib, 20nM) or DMSO control were added to the lineage depleted cells and refreshed every 24h. Media with mGM-CSF was refreshed every 2-3 days. In order to track differentiation, an aliquot of the cells was stained with anti-mouse CD11b efluor450 and Ly6G/Ly6C (Gr1) PE-Cy7 antibody (1:150, 15min, RT) on day 0, 2 and 7 of the experiment. Dead cells were excluded with 7-AAD.

##### 1.2 Human specimens

All biobanked specimens were evaluated by cytospin preparations, and only samples with a myelomonocytic cell/blast percentage >80% were included. Samples were collected at diagnosis of CMML or at refractory/progressed disease or secondary AML transformation as indicated in the Supplemental Table 1.

###### 1.2.1 Ex vivo cell death and cell cycle analysis of patient specimens

Human primary specimens were thawed and cultivated in SFEM II media (Stemcell Technologies, #09655), supplemented with 0.2% Antibiotic-Antimycotic (100X, Thermo Scientific), TPO (100ng/ml; Peprotech, #300-18), FLT3L (100ng/ml; Peprotech, #300-19),

SCF (100ng/ml; Peprotech, #300-07), IL-6 (100ng/ml; Peprotech, #200-06), IL-3 (100ng/ml; Peprotech, #200-03), UM729 (350nM; Stemcell Technologies, #72332), and SR1 (750nM) (Selleck Chemicals LLC, #S2858). Approximately $0.5\text{-}2x10^6$ cells were seeded in 1ml of media in a 24 well plate.

For cell death analysis, cells were incubated with either MEKi (Mirdametinib or Trametinib, 20nM, 24h) or DMSO vehicle control. Following the incubation period, cells were stained with panel 6 (Supplemental Table 7) (10min, RT). Following a washing step, cells were resuspended in Annexin binding buffer (BD Pharmingen) with the addition of Annexin V (1:20) and 7-AAD (1:20). Analysis of viable (Annexin $V^-$/7-$AAD^-$) cells was performed on a flow cytometer within 1h.

For cell cycle/proliferation analysis, cells were incubated with either MEKi (Mirdametinib or Trametinib, 20nM, 48h) or DMSO vehicle control. MEKi and DMSO vehicle were refreshed after 24h. Following the incubation period, each specimen was incubated with EdU (10µM, Click-iT™ Plus EdU Alexa Fluor™ 647 Flow Cytometry Assay Kit, Thermo Fisher Scientific, Waltham, MA, USA, #C10634). 1-2 hours later, cells were harvested and stained with Panel 6 (Supplemental Table 7). FxCycle Violet (1:1000 dilution, Thermo Fisher Scientific, #F10347) was added prior to acquisition with a flow cytometer.

##### 1.3 Patient-derived xenografts

Transplantation was performed as outlined in the main manuscript. Randomization was performed using the GraphPad QuickCalcs Web site: graphpad.com (accessed September 2025). Peripheral blood counts and engraftment checks were performed every 2 weeks of the treatment protocol. Engraftment of human CMML cells was monitored by regular blood sampling and flow cytometric analysis using Panel 4 (Supplemental Table 5). At experimental end point bone marrow and spleen cells were additionally stained for engraftment monitoring. Spleen weight, length and peripheral blood counts were measured at sacrifice.

In order to asses cell proliferation, single cell suspensions of bone marrow from Trametinib or vehicle treated PDX animals were resuspended in RPMI supplemented with 10% FCS and 1% Antibiotic-Antimycotic (100X, Thermo Scientific) in a 24 well plate at a cell density of $1x10^6$ per ml. Each specimen was incubated with EdU (10µM, Click-iT™ Plus EdU Alexa Fluor™ 647 Flow Cytometry Assay Kit, Thermo Fisher Scientific, Waltham, MA, USA, #C10634) in duplicates. 2 hours later, cells were harvested and stained with anti-human CD45 PE-Cy7 (Biolegend, Cat#982310; 1:100, 15min, RT). FxCycle Violet (1:1000 dilution, Thermo Fisher Scientific, #F10347) was added prior to acquisition with a flow cytometer.

##### 1.4 Flow Cytometry

Single cell suspensions of mouse tissue (prepared in 1.1.3) were stained for a myeloproliferative phenotype with antibodies listed in Supplemental Table 2 (15 min, RT). Enriched mouse hematopoietic cells, where lineage-committed cells were biotin-labelled, were incubated with streptavidin APC-Cy7 (20 µg/ml, BD, #554063, 15 min, 4°C). Next, cells were washed and HSC panel (Supplemental Table 3) or HSPC panel of antibodies was added to the cells (Supplemental Table 4) (15 min, RT). 7-AAD (1.5 µg/mL in PBS; BD) addition was used to exclude dead cells during flow cytometry.

To analyze human engraftment in the PDX model, single cell suspensions of bone marrow and peripheral blood were blocked with a mix of human (Human TruStain FcX, Biolegend, #422302) and mouse (FcR Blocking Reagent, mouse, Miltenyi Biotec, 130-092-575) FcX block (1:20, 10 min, 4 °C). Next the engraftment panel of antibodies (Supplemental Table 5) was added (30 min, 4 °C). Following the incubation period, cells were washed, resuspended in Sytox Red Dead Cell Stain (1:1000, Invitrogen) and measured immediately. Cells were acquired on

CytoFLEX LX or CytoFLEX S (Beckman Coulter, Krefeld, Germany) with CytExpert software. Flow cytometry data were analyzed using FlowJo software (V10.10, TreeStar).

###### 1.4.1. Phospho-Flow

The phospho-flow protocol was adapted from Kalaitzidis et al[6]. In brief, lineage depleted bone marrow cells were seeded in a 24-well plate in a cell density of 0.5-1.5 x $10^6$ per well in starvation media (IMDM + 2% FCS) for 1h. Next cells were washed with PBS + 1% BSA and labelled with Streptavidin APC-Cy7 (15 min, 4°C). After this, cells were first stained with a panel of antibodies (Supplemental Table 6) including Fixable Viability dye to exclude dead cells (30min, 4°C). Following a washing step, growth factors or vehicle were added to the cell suspensions SCF (100ng/ml; Peprotech, # 250-03), Flt3L (50ng/ml; Peprotech, # 250-31L), GM-CSF (20ng/ml; Peprotech, # 315-03) for 10min at 37 degrees. Cells were fixed with paraformaldehyde (1.5%, 10 min, RT) and after washing (PBS + 0.50% BSA + 0.02% NaN3), permeabilized with ice-cold acetone. Finally intracellular antibodies against p-Erk (Cell signaling Technology, Cat. #4374), p-Akt (Cell signaling Technology, Cat. #2336) or p-S6 (Cell signaling Technology, Cat. #4803) conjugated with Alexa Fluor 488 were added to the cells (20 min, RT) and acquired on a flow cytometer.

##### 1.5 Immunoblot

Ice-cold RIPA-Buffer (Sigma), supplemented with protease and phosphatase inhibitor cocktails (Thermo Fisher Scientific), was added to isolated total bone marrow cell pellets to obtain cell lysis. Protein concentration was measured using the DC Protein Assay kit (Bio-Rad, Hercules, CA, USA) following the manufacturer’s protocol. Immunoblots were then performed as previously described [7, 8] using Mini-PROTEAN TGX gels for electrophoresis (Bio-Rad) and the Bio-Rad Trans Blot TurboBlotting System for transfer. Polyvinylidene difluoride membranes (Bio-Rad) were incubated with anti-EZH2 (#3147S; Cell Signaling Technologies, Danvers, MA, USA), anti-ERK (#M5670; Sigma-Aldrich, St. Louis, MO, USA), anti-pERK (#4370; Cell Signaling), anti-H3K27me3 (#9733; Cell Signaling) and anti-Vinculin (#ab129002; Abcam, Cambridge, UK). The intensity of the bands was compared using ImageJ.

##### 1.6 Immunohistochemistry

Mouse spleen tissue samples were fixed in formalin, dehydrated and embedded in paraffin. Sectioned slides were rehydrated and followed by standard H&E staining protocol. PCNA immunostaining was performed on rehydrated and blocked spleen sections. Sections were then incubated with anti-PCNA (1:500, Merck, #MAB424) for 60 min, washed and visualized with the help of AEC Substrate Chromogen (Abcam, #AB103742). Area covered by PCNA positive cells was quantified with Image J Color deconvolution and normalized to area covered by total cell infiltrate.

For pMEK IHC sections were deparaffinized, blocked and stained with anti-pMEK (1:50; Cell Signaling, #CS9121). A scoring system for pMEK IHC was adapted from Ormanns et al[9, 10] where the nuclear and cytoplasmic staining intensity (0 - no, 1 - weak, 2 - moderate, and 3 – strong staining) were added to the score for the percentage of positive cells (0 - negative, 1 - less than 25 %, 2 – 25 % to 50 %, 3 - more than 50 % positive staining cells). Finally, nuclear and cytoplasmic score were summarized. Specimens were defined as pMEKhigh if the total score was 6 or higher. Patient data for samples used for p-MEK scoring can be found in Supplemental Table 1.

##### 1.7 Database retrieval and statistical analysis

Database retrieval was performed as outlined in the main manuscript. For AML ex-vivo drug sensitivity data,[1] $EZH2^{inact}$ was defined as the presence of homozygous or heterozygous EZH2 mutation(s), deletion of chromosome 7 or 7q, and/or reduced EZH2 expression. As only transcriptomic were available, we defined reduced EZH2 expression as an RNAseq-derived z-score $\leq -2$ relative to the cohort mean, following the default cBioPortal threshold for expression outliers. Cases with incomplete mutational or drug sensitivity data, non-CMML/AML diagnoses, or unclear EZH2 status (e.g., derivative chromosome 7) were excluded. Gene sets of canonical pathways (BIOCARTA_ERK_PATHWAY, BIOCARTA_AKT_PATHWAY and BIOCARTA_p38/MAPK_PATHWAY) were downloaded from https://www.gsea-msigdb.org/gsea/index.jsp.[2, 3] The z-scores for gene expression of individual genes in the pathways were downloaded for the samples that had included drug sensitivity data and the median z-score was taken for analysis.
